# Pannexin 3 channels regulate architecture, adhesion, barrier function and inflammation in the skin

**DOI:** 10.1101/2022.07.13.499920

**Authors:** Brooke L. O’Donnell, Rafael E. Sanchez-Pupo, Samar Sayedyahossein, Mehdi Karimi, Mehrnoosh Bahmani, Christopher Zhang, Danielle Johnston, John J. Kelly, C. Brent Wakefield, Kevin Barr, Lina Dagnino, Silvia Penuela

## Abstract

The channel-forming glycoprotein Pannexin 3 (PANX3) functions in cutaneous wound healing and keratinocyte differentiation, but its role in skin homeostasis through aging is not yet understood. We found that PANX3 is absent in newborn skin but becomes upregulated with age. We characterized the skin of global *Panx3* knockout mice (KO) and found that KO dorsal skin showed sex-differences at different ages, but generally had reduced dermal and hypodermal areas compared to aged-matched controls. Transcriptomic analysis of KO epidermis revealed reduced E-cadherin stabilization and Wnt signaling compared to WT, consistent with the inability of primary KO keratinocytes to adhere in culture, and diminished epidermal barrier function in KO mice. We also observed increased inflammatory signaling in KO epidermis and higher incidence of dermatitis in aged KO mice compared to wildtype controls. These findings suggest that during skin aging, PANX3 is critical in the maintenance of dorsal skin architecture, keratinocyte cell-cell and cell-matrix adhesion and inflammatory skin responses.

## INTRODUCTION

Pannexins (PANX1, PANX2, PANX3) are a family of channel-forming glycoproteins (Panchin et al., 2000) whose primary function is to form large-pore, single-membrane channels to allow the passage of ions and metabolites during autocrine and paracrine signaling (Penuela et al., 2013) All three pannexins are found in the skin (Le Vasseur et al., 2014, Penuela et al., 2007).

To date, PANX3 has been reported in cartilage (Iwamoto et al., 2010), teeth (Fu et al., 2015), bone (Ishikawa et al., 2011), skeletal muscle (Langlois et al., 2014), skin (Penuela et al., 2007), adipose (Pillon et al., 2014), mammary glands, the small intestine (Bond et al., 2011), a subset of blood vessels (Lohman et al., 2012), and the male reproductive tract (Turmel et al., 2011). In chondrocytes and osteoblasts, PANX3 is induced during differentiation, functioning at the cell surface as an ATP conduit and/or in the endoplasmic reticulum (ER) as a Ca^2+^ leak channel (Ishikawa et al., 2011, Iwamoto et al., 2010).

Our group previously demonstrated that PANX1 is highly expressed in young mouse dorsal skin, but becomes downregulated with age. Moreover, when comparing the dorsal skin architecture of wildtype (WT) and global *Panx1* knockout (KO) mice, we discovered that *Panx1* KO dorsal skin has a significantly reduced dermal and increased hypodermal area as well as compromised wound healing abilities compared to age-matched controls (Penuela et al., 2014). However, the role of PANX3 in skin homeostasis during aging is unknown.

PANX3 protein was found to be present in all epidermal layers of both thin and thick skin of 3-week-old C57BL/6 mice as well as sebocytes with a predominantly intracellular localization (Celetti et al., 2010). Similarly in aged human skin, PANX3 exhibits an intracellular localization pattern in all layers of the epidermis (Penuela et al., 2007) and in supporting structures such as sebaceous glands, eccrine glands, hair follicles and blood vessels (Cowan et al., 2012). PANX3 can be detected in the epidermis as early as E13.5 (Celetti et al., 2010), but there are conflicting results on its expression past birth (Celetti et al., 2010, Ishikawa et al., 2019). Thus, the regulation of PANX3 levels with skin aging remains to be determined.

Global *Panx3* KO mice were shown to have a reduced epidermal and dermal thickness at P4, but these phenotypes were no longer present at P10, P20 and P38, and P4 *Panx3* KO skin also showed marked reduction in keratinocyte differentiation markers. Furthermore, when PANX3 was overexpressed in the HaCaT human keratinocytic cell line, PANX3 increased keratinocyte differentiation, inducing the expression of key drivers of differentiation such as Epiprofin and Notch1. At the same time ectopic PANX3 expression reduced proliferation by promoting cell arrest in the G_0_/G_1_ phase through protein kinase B/nuclear factor of activated T-cells signaling triggered by ATP and Ca^2+^ release through PANX3 channels (Zhang et al., 2021). Despite these findings, the effect of *Panx3* ablation on the architecture of aged skin and the expression and function of endogenous PANX3 levels in keratinocytes has yet to be investigated.

Zhang *et al*. (2019) also demonstrated that *Panx3* KO mice have delayed cutaneous wound healing capacities compared to controls. PANX3 was particularly important during re-epithelialization, where keratinocytes proliferate, migrate, and differentiate to replace the lost epithelium via epithelial-to-mesenchymal transition (EMT). Specifically, during transforming growth factor β (TGF-β) stimulated-EMT, PANX3 was found to act as both a cell surface and ER channel for the passage of ATP and Ca^2+^, respectively, in PANX3-overexpressing HaCaT keratinocytes (Zhang et al., 2019). However, the function of PANX3 in skin homeostasis and aging in the absence of challenges such as wound healing has yet to be fully investigated.

Using the global *Panx3* KO mouse model previously developed by our group (Moon et al., 2015), we investigated PANX3 expression and function in the maintenance of proper skin architecture with aging. We found that PANX3 is highly expressed in primary keratinocytes where it is critical for keratinocyte adhesion. PANX3 levels increase with aging and its presence is required to maintain skin structure, epidermal barrier function and inflammatory responses, with *Panx3* KO epidermis exhibiting impaired Wnt and E-cadherin signaling, a compromised barrier and increased incidence of dermatitis.

## RESULTS

### PANX3 is upregulated in WT male and female neonatal dorsal skin

To determine the regulation of PANX3 levels throughout dorsal skin aging, both male and female (Figure 1a,b) WT dorsal skin were collected from mice aged P0 to 1 year. Both sexes exhibited the same PANX3 expression pattern, with PANX3 undetectable in newborn skin, becoming upregulated at P4 and remaining at high levels with aging. When directly comparing male and female WT dorsal skin, we found that the PANX3 protein levels detectable in P4 (Figure 1c) and 22-week-old (Figure S2a) mice showed no differences between the sexes, consistent with similar *Panx3* mRNA levels in P4 dorsal skin (Figure 1d). Due to N-glycosylation, PANX3 bands can appear as a singlet or doublet in Western blots. In doublets, the lower species correspond to the high mannose glycosylated form Gly1 and the higher species corresponds to the complex glycoprotein Gly2 (Penuela et al., 2009). In the dorsal skin, Gly1 was present, consistent with previous research which indicates a predominantly intracellular PANX3 localization in the epidermis and adnexal skin structures (Celetti et al., 2010, Cowan et al., 2012). Furthermore, we found a gradual increase in PANX3 levels from P0 to P4 (Figure 1e), and PANX3 was detected in dorsal skin as late as 18 months of age at comparable levels in both male and female WT mice (Figure S2b; full blots Figure S1a-d).

**Figure 1.**
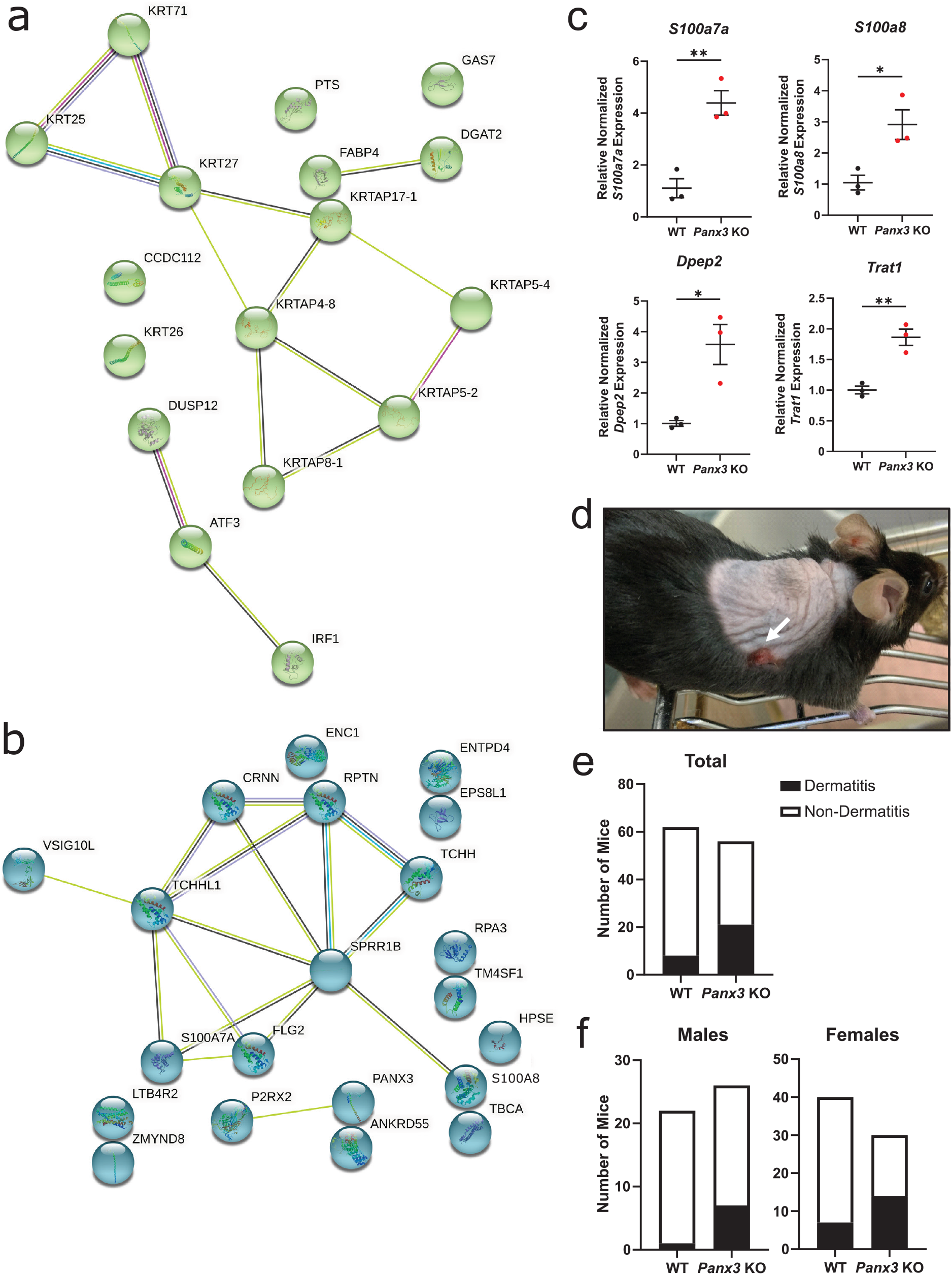
PANX3 is expressed in dorsal skin beginning at P2, with the same pattern seen in both sexes. An immunoblot showing PANX3 in male (**a**) and female (**b**) WT mouse dorsal skin from P4 to 1 year of age, with a significant increase in PANX3 levels from P0 to P4, but no differences from P4 to 1 year (males *N=3*, females *N=4*, \*\**p*<0.01, *** *p*<0.001, \*\*\*\**p*<0.0001 compared to WT P0). KO dorsal skin as negative controls. (**c**) PANX3 levels do not differ between male and female P4 WT dorsal skin (NS, *p*>0.05). (**d**) In female WT mice, PANX3 levels gradually increase from P0 to P4 (*N*=4, \*\**p*<0.01, *** *p*<0.001, \*\*\*\**p*<0.0001). Human embryonic kidney 293 cells ectopically expressing a mouse PANX3 plasmid (HEK+PANX3) were used as a positive control. PANX3 levels normalized to glyceraldehyde-3-phosphate dehydrogenase (GAPDH; protein loading control) and compared using an unpaired t-test or a one-way ordinary measures ANOVA with Tukey’s multiple comparisons test. Protein sizes in kDa, error bars ±SEM.

We also compared PANX1 and PANX2 levels (Figure S3) in WT and *Panx3* KO dorsal skin and found that only PANX2 isoform 202 protein levels differed, with a significant reduction in KO skin, showing no indication of compensation from other family members with *Panx3* ablation. These findings suggest, that although absent in newborns, PANX3 is present in neonatal dorsal skin and its expression persists with skin aging.

### *Panx3* KO mice have reduced dorsal skin thickness at different ages in males and females

Upon closer histological examination of global *Panx3* KO dorsal skin we identified sex-specific architectural differences compared to WT controls. In male dorsal skin (Figure 2), the hypodermal area of KO mice at 4 weeks was significantly reduced. Additionally, at 1 year, KO males showed both a reduced epidermal and dermal area, and hypodermal area. However, KO female dorsal skin (Figure 3) had significantly reduced hypodermal areas from P0 to 1 year, but a reduced epidermal and dermal area was only observed in P0 and 1-year-old mice. Contrarily, there were no significant differences in epidermal areas of both male and female paw skin regardless of genotype or mechanical challenge (Figure S4,5). Taken together, these results demonstrate PANX3 is required for the maintenance of proper dorsal skin architecture, particularly in aged skin.

**Figure 2.**
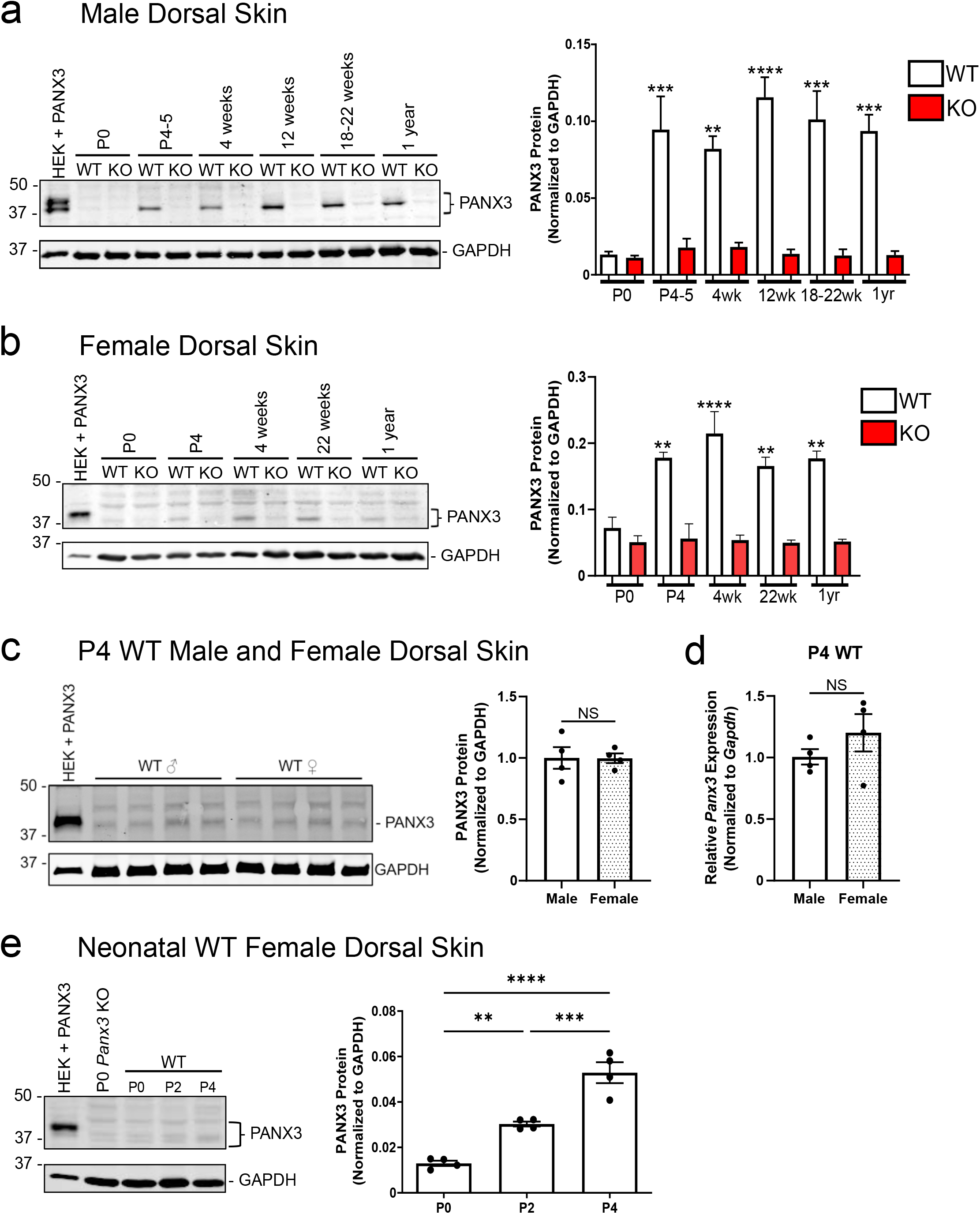
In male KO dorsal skin, the epidermal and dermal and hypodermal areas are decreased in aged mice. Histological analysis of male murine dorsal skin showed the hypodermal area at 4 weeks (\*\*\*\**p*<0.0001) and 1 year (\*\*\**p*<0.001) was significantly decreased in KO mice compared to controls (unpaired t-test). The epidermal and dermal area of 1-year-old KO mice was also reduced (\*\**p*<0.01). No significant differences were seen in any other measurements (*p*>0.05). Upper brackets represent epidermal and dermal layers (minus the *stratum corneum*), lower brackets represent the hypodermis. *N=5, n*=15. Error bars ±SEM. Scale bars: 50μm at P0, 100μm at P4, 4 weeks, 22 weeks and 1 year. NS, no significance.

**Figure 3.**
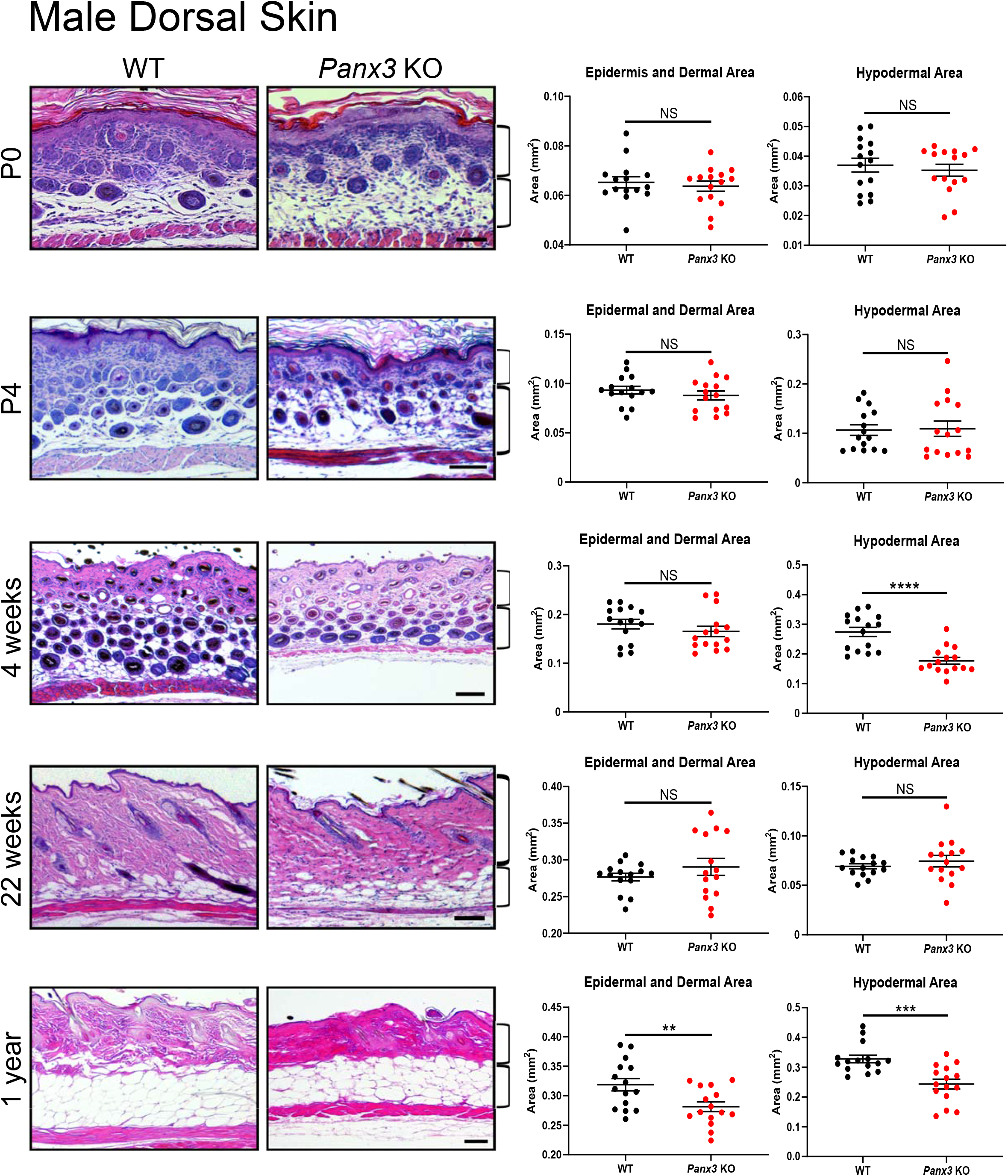
In female KO dorsal skin, the hypodermal area is reduced at all ages. Histological analysis of female murine dorsal skin showed a decreased hypodermal area from P0 to 1 year in KO mice (\**p*<0.05, \*\**p*<0.01, \*\*\**p*<0.001) \*\*\*\**p*<0.0001; unpaired t-test). Reduced epidermal and dermal areas were seen in P0 and 1-year-old KO dorsal skin. No significant differences were seen in any other measurements (*p*>0.05). *N*=5, *n*=15. Upper brackets represent epidermal and dermal layers (minus the *stratum corneum*) and lower brackets represent the hypodermis. Error bars ±SEM. Scale bars: 50μm at P0, 100μm at P4, 4 weeks, 22 weeks and 1 year. NS, no significance.

### *Panx3* KO neonates have compromised epidermal barrier function

Although the dermal area is reduced (Figure S6a), we focused our investigation on the epidermis of *Panx3* KO skin since we could not detect PANX3 protein in isolated cultured dermal fibroblasts, *Panx3* transcripts remained unchanged with TGF-β-induced fibroblast activation and *Panx3* KO contains similar collagen levels to WT dorsal skin (Figure S6b-d). Additionally, PANX3 function has been investigated in keratinocyte overexpression systems, but no studies have explored the effects of endogenous PANX3 in keratinocytes (Celetti et al., 2010, Zhang et al., 2021, Zhang et al., 2019). Thus, we isolated the epidermis of WT and KO mice to investigate endogenous PANX3 expression and the effect of *Panx3* ablation in these cells. We identified that *Panx3* transcripts were highly expressed in the WT neonatal epidermis, but histological measurements showed no differences in the epidermal area between WT and KO dorsal skin (Figure 4a,b).

**Figure 4.**
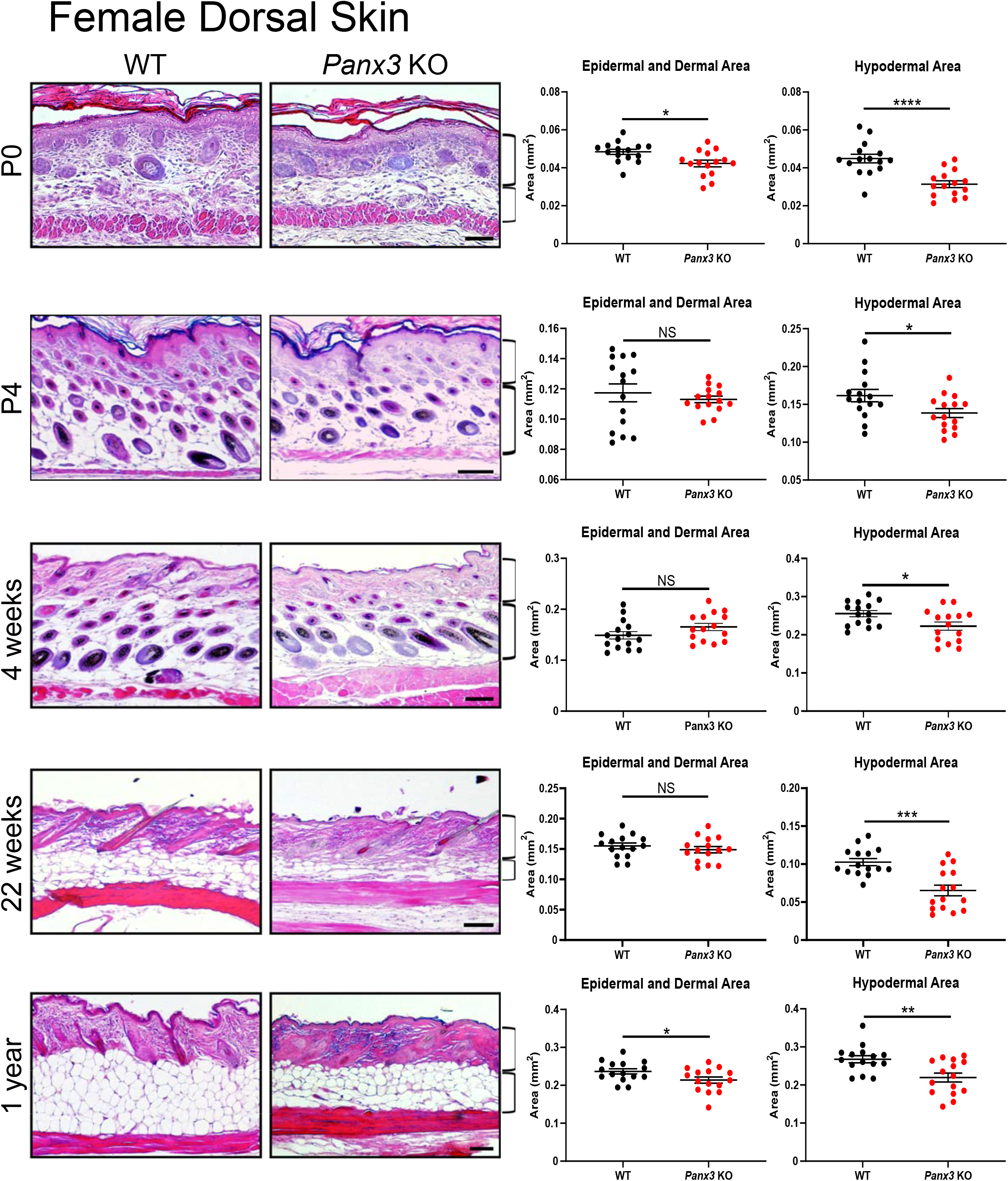
Epidermal barrier function is compromised in neonatal KO dorsal skin. (**a**) RT-qPCR showed *Panx3* transcript is present in WT P4 neonatal epidermis (*N*=3, pooled samples of 4-6 male and female pups). *Gapdh* was used to calculate normalized mRNA expression via ΔΔCT. (**b**) Histological analysis showed **e**pidermal area measurements from P0 and P4 dorsal skin (*N*=5, n=15) were not significantly different (NS, no significance; unpaired t-test). (**c**) P2 pups were subjected to toluidine blue staining to test epidermal barrier function (*N*=15). Dye infiltration seen in all KO pups, but not in unstained controls or WT pups, indicates compromised barrier function. Intentional barrier disruption using a scalpel (Cut). Arrows point to areas of increased dye infiltration KO mice and corresponding areas in WT pups for comparison.

One of the main functions of the epidermis is its action as an protective barrier against pathogens and water loss (Baroni et al., 2012), and to investigate the epidermal barrier function in KO mice we performed a toluidine blue dye assay. All male and female KO pups had blue staining on all areas of the skin, indicating dye permeation, not seen in WTs or unstained controls. Additionally, prominent blue staining was observed in the mouth/nose area, paws and tails of KO mice (Figure 4c). Ultimately, these findings uncovered a novel phenotype of *Panx3* KO mice, demonstrating that KO skin exhibits a leaky epidermal barrier and reduced keratinocyte cell-cell adhesion.

### *Panx3* KO neonatal epidermis has reduced adhesion signaling

To understand the signaling mechanisms behind the compromised epidermal barrier upon *Panx3* deletion, we performed a Clariom™ S transcriptomic profiling analysis of WT and KO neonatal epidermis (Figure S8). Of the 22,206 genes tested, 999 differentially expressed genes (DEGs) with a fold change of ±1.5 were identified, including 539 upregulated and 460 downregulated genes in KO neonatal epidermis. When DEGs were organized into functional categories and signaling pathways relevant in skin health and conditions such as psoriasis, dermatitis, and skin cancer (Figure S7d), we found that many gene candidates were implicated in skin development and barrier function, consistent with our findings of altered skin structure and barrier integrity. Using Gene Set Enrichment Analysis (GSEA), gene sets for E-cadherin stabilization and Wnt signaling (Figure 5a) were the most negatively enriched in KO neonatal epidermis. Decreased cadherin 1 (*Cdh1*), vinculin (*Vcl*), Wnt family member 3 (*Wnt3*) and low-density lipoprotein receptor-related protein 5 (*Lrp5*) transcripts were seen in KO neonatal epidermis (Figure 5b).

**Figure 5.**
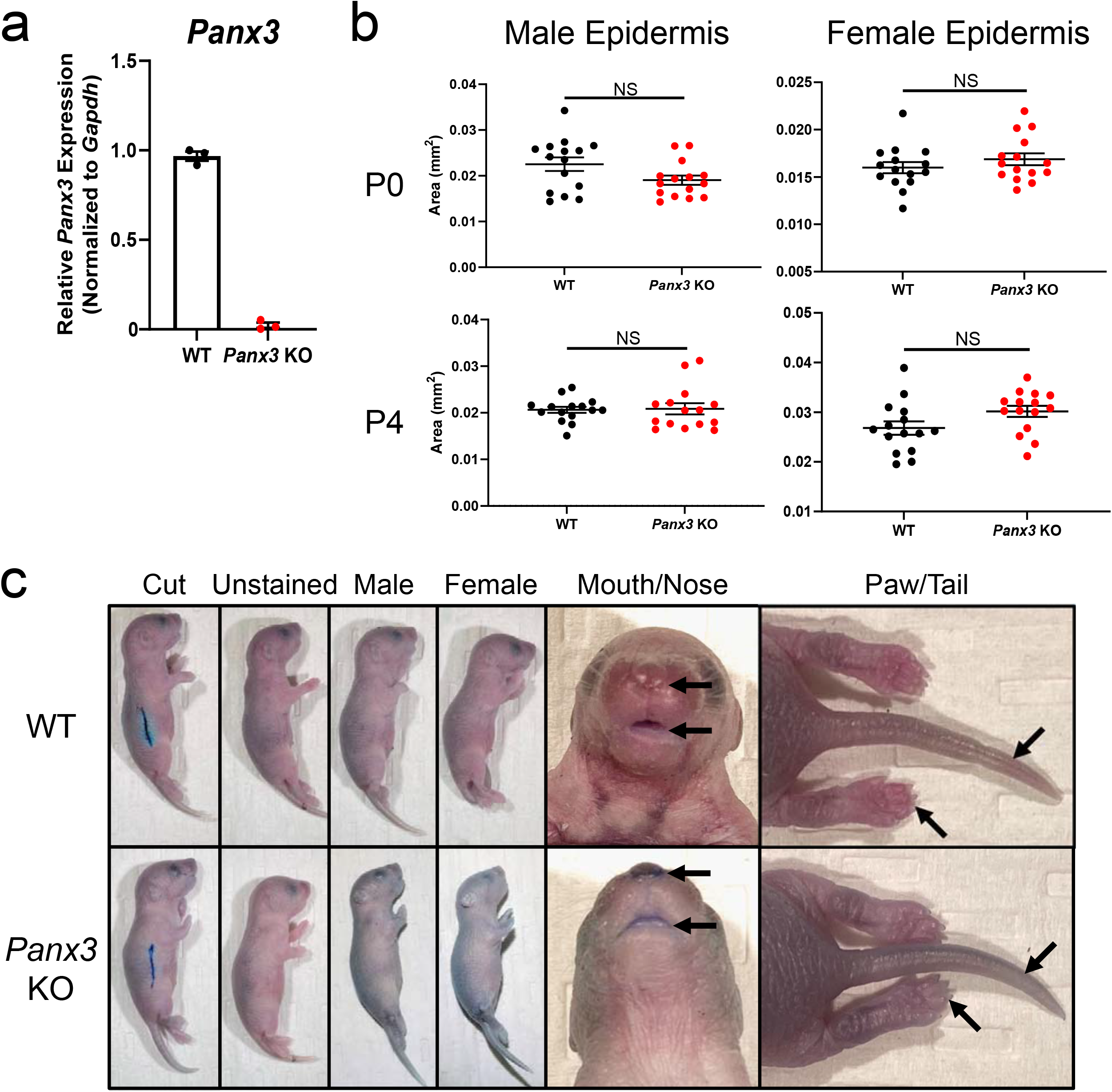
Adhesion signaling is reduced in KO neonatal epidermis. (**a**) E-cadherin stabilization and Wnt signaling were the most negatively enriched gene sets in KO neonatal epidermis (ES<0) via GSEA pathway analysis (*N*=3). (**b**) Vinculin (*Vcl*), Wnt family member 3 (*Wnt3*) and low-density lipoprotein receptor-related protein 5 (*Lrp5*, \*\**p*<0.01) were significantly reduced and cadherin 1 (*Cdh1, p*=0.08) trended lower in KO neonatal epidermis. Unpaired t-test. *Gapdh* was used for normalization. Bars: ±SEM. (**c**) Brightfield images of WT (day 7) and KO (day 3) primary keratinocytes on untreated tissue culture plastic (*N*=18, 9 per sex). (**d**) Live and dead cell percentages of WT and KO neonatal epidermis before plating (*N*=9, 5 males, 4 females) (*p*>0.05, NS, no significance, unpaired t-test) between genotypes. (**e**) Brightfield images of KO keratinocytes plated on tissue culture plastic coated with 50μg/mL collagen, 100μg/mL collagen, laminin or fibronectin (*N*=6, 3 per sex). Scale bar: 50μm.

Even though keratinization was identified as the most positively enriched pathway in GSEA and Reactome analyses, most differentially expressed genes in KO neonatal epidermis were critical in epidermal barrier function rather than differentiation. Additionally, no differences were seen in differentiation markers in the neonatal epidermis of each genotype, and *Panx3* transcript levels did not change upon Ca^2+^ differentiation of isolated WT primary keratinocytes (Figure S9).

### KO keratinocytes do not adhere in culture

To explore whether the negatively enriched adhesion signaling was reflected in cultured keratinocytes, we isolated WT and KO keratinocytes from P4 dorsal skin (Figure S8a). WT keratinocytes adhered to the culture dish and spread out to make contacts with neighboring cells to form clusters. In contrast, KO keratinocytes did not adhere to tissue culture plastic or contact neighboring cells, remaining ‘balled up’ in structure and lifting off the plate shortly after isolation (Figure 5c). However, this phenotype was not due to reduced KO keratinocyte viability, since there were no significant differences in live and dead cell percentages between the neonatal epidermis of each genotype immediately before cell plating (Figure 5d). It was also independent of the extracellular matrix protein used for culturing, since KO keratinocytes exhibited the same low adherence when plated onto surface-treated plates coated with 50μg/mL collagen, 100μg/mL collagen, laminin or fibronectin (Figure 5e). Interestingly, isolated dermal fibroblasts from each genotype could be cultured and passaged with no signs of adherence or growth deficits, indicating this KO phenotype is cell-type specific. Ultimately, these results illustrate that KO keratinocytes show both cell-cell and cell-matrix adhesion deficits, reflected in their reduced E-cadherin and Wnt signaling, inability to adhere in culture and the compromised epidermal barrier.

### Aged *Panx3* KO mice have increased risk of dermatitis

A STRING analysis produced two clusters related to inflammation, with one plot associated with the formation of the cornified envelope and *Staphylococcus aureus* infection (Figure 6a), and another cluster associated with the cornified envelope and S-100 type calcium binding proteins (Figure 6b)—which are often upregulated in psoriatic and dermatitis inflammatory lesions (Eckert et al., 2004). Most inflammatory gene candidates within these clusters, such as S100 calcium binding protein A7A (*S100a7a*) and A8 (*S100a8*), dipeptidase 2 (*Dpep2*) and T cell receptor associated transmembrane adaptor 1 (*Trat1*), were significantly increased in KO neonatal epidermis (Figure 6c).

**Figure 6.**
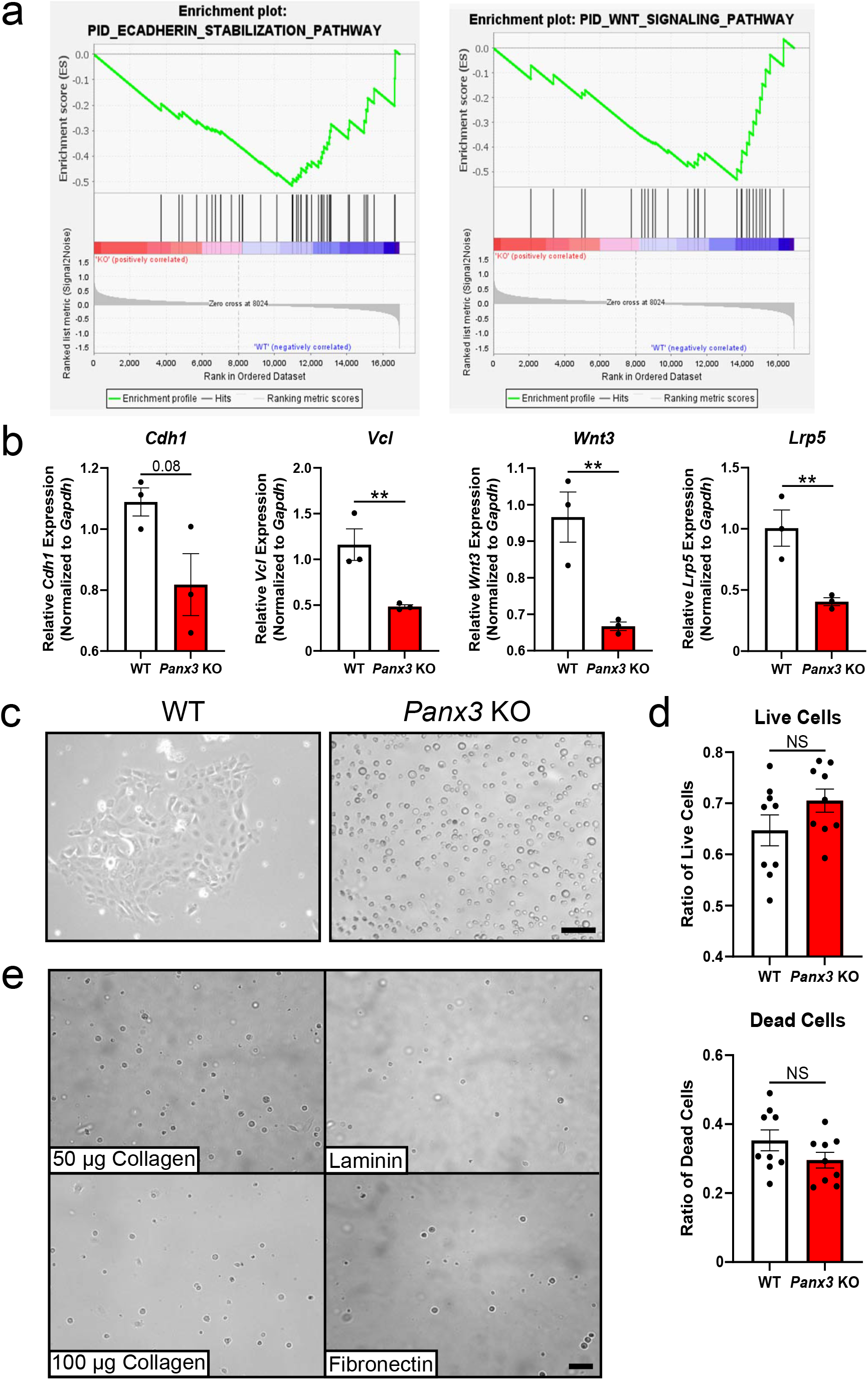
Inflammation signaling and dermatitis in KO mice. STRING plots were generated using the 100 highest ranked differentially expressed genes showing ±2-fold changes. Cluster (**a**) was associated with cornification, formation of the cornified envelope and *Staphylococcus aureus* infection. Cluster (**b**) was associated with the cornified envelope and S-100/ICaBP type calcium binding domain. (**c**) S100 calcium binding protein A7A (*S100a7a*) and A8 (*S100a8*), dipeptidase 2 (*Dpep2*) and T cell receptor associated transmembrane adaptor 1 (*Trat1*) transcript levels were significantly increased in KO neonatal epidermis (\**p*<0.05, \*\**p*<0.01, unpaired t test). Bars represent ±SEM. (**d**) WT female with dermatitis (53% score), lesion denoted by white arrow. (**e**) Dermatitis incidence was analyzed in a cohort of 118 aging WT and KO mice with an association between genotype and dermatitis (*p*<0.01). (**f**) Dermatitis incidence by sex. Dermatitis and genotype were only linked in females (males *p*=0.055; females *p*=0.017). Fisher’s exact test.

Since dermatitis is common in C57BL/6 mice (Hampton et al., 2012), and a compromised epidermal barrier increases the risk of skin infection and dermatitis (Bozek et al., 2020, Jacoby et al., 2002), we investigated whether the increased inflammatory signaling was reflected in the incidence of dermatitis in our aging mouse colony where we found PANX3 is still present in aged dorsal skin (Figure S1b). Figure 6d shows an example of a WT mouse diagnosed with moderate dermatitis, where in this typical case the small lesion appeared red and inflamed with some scabbing. In total, 8/62 WT and 20/56 KO mice developed dermatitis (Figure 6e) and we found an association between genotype and dermatitis incidence (*p*<0.01), with KO mice having a 4.05 times higher risk of developing dermatitis than controls. However, this risk differed depending on the sex of the mouse since most dermatitis cases were in female mice, with 7/40 WT and 14/30 KO females developing dermatitis as opposed to only 1/22 WT and 7/26 KO males (Figure 6f). There was an association between genotype and dermatitis diagnosis only in the female cohort (*p*<0.05); however, the male cohort was close to significance (*p*=0.055). The odds ratios also showed sex differences, with KO females at a 4.13 times greater risk of being diagnosed with dermatitis than WT females, whereas KO males were at a 7.74 times greater risk. Thus, it seems the compromised epidermal barrier function seen in KO mice persists with age but there may be sex differences.

## DISCUSSION

In this study, we found that PANX3 is a key regulator of skin homeostasis throughout aging, regulating keratinocyte adhesion, inflammation, and epidermal barrier integrity. When investigating the mechanism behind these functions through an expression profiling approach, we determined the phenotypes observed in the *Panx3* KO epidermis most likely result from deficiencies in Wnt signaling and E-cadherin stabilization. Wnt signaling plays an integral role in keratinocyte proliferation and differentiation, hair follicle development and cutaneous wound response (Lang et al., 2019). Previous research using a keratinocyte-specific KO of wntless Wnt ligand secretion mediator illustrated that abrogating Wnt secretion altered keratinocyte differentiation and impaired the epidermal barrier (Augustin et al., 2013). Thus, we speculate that the reduced Wnt signaling in KO neonatal epidermis contributes to perturbations in epidermal barrier function. Using the punch biopsy model, Zhang *et al*. (2019) found that at the wound site, β-catenin mRNA was significantly reduced in KO wounded skin compared to controls. Recently, we discovered that PANX1 binds directly to β-catenin, a main player in the Wnt pathway, for signaling purposes most likely independent of PANX1 channel function (Sayedyahossein et al., 2021). Thus, it is possible PANX3 may also act as a scaffold protein to β-catenin or other Wnt signaling components, functioning through its interactome rather than as a channel. Since PANX3 can act as an ER Ca^2+^ leak channel in keratinocytes (Zhang et al., 2019), it is also possible that the lack of PANX3 disrupts intracellular calcium signaling and thus calcium-dependent adhesion, by preventing the downstream phosphorylation of α-, β- and p120 catenins and their subsequent binding and stabilization of the adherens junction complex (Bikle et al., 2012). Ultimately, resulting in the compromised barrier function exhibited in KO neonates.

The reduced epidermal and dermal area observed parallels the reductions in epidermal and dermal thickness observed in P4 KO skin of another global *Panx3* KO mouse model. In this alternate model, the phenotypes seem to be lost by P10, similar to our own findings in which the reduced epidermal and dermal area was only present in P0 females and 1-year-old KO mice. Additionally, the representative histological images of P4 and P38 KO mice illustrate a markedly reduced subcutaneous fat thickness, consistent with our findings in female KO mice; however, the sexes of the mice were not reported (Zhang et al., 2021).

Since all three pannexins are expressed in the skin, there is a chance of compensation from other pannexin family members when one is ablated. Notably, PANX3 levels were significantly increased in *Panx1* KO dorsal skin (Penuela et al., 2014). However, the reciprocal cannot be said for PANX1 in *Panx3* KO mice, since we did not see any significant difference in PANX1 levels compared to controls, but this does not rule out the possibility of functional compensation. Phenotypically, both *Panx1* and *Panx3* KO models exhibit a reduction in the epidermal and dermal area, but unlike *Panx1* KO mice which exhibit increased subcutaneous fat, the hypodermal area is significantly reduced in the *Panx3* KO mice (Penuela et al., 2014). We previously found that the reduction in hypodermal area in the *Panx3* KO was due to reduced adipocyte cell number rather cell size (Wakefield et al., 2021). PANX2 isoform 202 was downregulated in *Panx3* KO dorsal skin compared to controls which is, to our knowledge, the first report of a reduction in pannexin levels when another family member is ablated.

Despite the increased numbers of dermatitis in female mice, the odds ratio of developing dermatitis was much higher in male KO mice than in females, indicating PANX3 plays a clear role in regulating inflammatory responses in both sexes, but the effect is more prominent in males. The overall increased dermatitis incidence in females may have occurred due to multiple factors, such as increased incidences of dermatitis secondary to barbering and ulcerative dermatitis in female mice (Garner et al., 2004, Hampton et al., 2012), and the 46% higher sample size of females in our study. These suggest higher numbers of dermatitis diagnosis in females regardless of genotype. However, a deletion of *Panx3* clearly increases the incidence of dermatitis in aged mice, suggesting a role in inflammation. Previous findings have been contradictory on the role of PANX3 in inflammation, suggesting PANX3 action may be tissuespecific. In healthy, unchallenged dorsal skin, we found that PANX3 seems to play more of an anti-inflammatory role evidenced by its ablation increasing inflammatory signaling. Using the same KO mouse model we previously showed that *Panx3* deletion reduced the inflammatory index in male visceral fat (Wakefield et al., 2021). Despite these differences, the contrary findings may not result from tissue dependence of PANX3 function, but rather age differences since visceral fat findings were in 30-week-old mice, whereas dermatitis cases were documented in 10.5-18-month-old mice. This is not the first instance of age-related effects of PANX3 since our group previously demonstrated the role of PANX3 in age-related acceleration of osteoarthritis (Moon et al., 2015), which contrasted the protective effects of injury-induced osteoarthritis in young mice (Moon et al., 2021).

Collectively, through the characterization of our global *Panx3* KO mouse, we discovered that PANX3 plays an integral role in the structure and function of dorsal skin, inflammatory responses and barrier function throughout aging and is requiredfor keratinocyte adhesion. By understanding the role that PANX3 plays in maintaining healthy skin, we will be better equipped to identify its dysfunction in inflammatory skin conditions such as psoriasis or dermatitis, or other skin pathologies such as keratinocytic skin cancers in which PANX3 is downregulated (Cowan et al., 2012, Halliwill et al., 2016) and may act as a tumor suppressor in malignant transformation of the skin.

## MATERIALS AND METHODS (Detailed methods in supplementary section)

### Animals and animal ethics

Global *Panx3* KO mice used were previously generated by our group and congenic C57BL/6N WT mice were used as controls (Moon et al., 2015). Animal experiments followed guidelines and protocols approved by the Animal Care Committee at the University of Western Ontario (#2019-069).

### Protein extraction and western blotting

Western blots of dorsal skin tissue were performed as reported (Abitbol et al., 2019), using 40μg per sample. Primary antibodies included: anti-PANX1 CT-395 0.5μg/mL, anti-PANX2 CT-523 4μg/mL and anti-PANX3 CT-379 1μg/mL (Penuela et al., 2009); generated anti-PANX3 CT-379-M1 and M2 unpurified supernatant 1:100, and anti-GAPDH 0.2μg/mL (Millipore Sigma, Burlington, MA).

### Generation of PANX3-specific antibodies

Carboxyl-terminal amino acid residues 379-392 (KPKHLTQHTYDEHA) of mouse PANX3 were used in mouse monoclonal antibody generation by Genemed Biotechnologies, Inc. (Torrance, CA). Unpurified hybridoma supernatant from two clones with the highest specificity and signal intensity were selected and designated PANX3 CT-379-M1 and PANX CT-379-M2.

### Histology

Histological staining and measurements were performed as described previously (Abitbol et al., 2019).

### Epidermal barrier function assay

Epidermal barrier function of P2 WT and KO pups was assessed using toluidine blue staining as reported (Press et al., 2017).

### Clariom™ S expression profiling and analysis

RNA from WT and KO neonatal epidermis were sent for Clariom™ S, Mouse expression profiling. Profiling data was analyzed using TAC (ThermoFisher Scientific, Rockford, IL) and RStudio Cloud (RStudio Team, 2022) software to find DEGs with a gene-level fold change of minimum ±1.5 and p<0.05. Further pathway analysis was conducted using GSEA (Mootha et al., 2003, Subramanian et al., 2005), Reactome (Fabregat et al., 2016, Fabregat et al., 2017) and STRING (Szklarczyk et al., 2019, Szklarczyk et al., 2021) databases.

### RNA isolation and RT-qPCR

RNA isolation and RT-qPCR were performed as described by Abitbol et al. (2019). Transcript abundance was normalized to *Gapdh*, presented as relative to one WT mean value and analyzed using the ΔΔCT method in Excel (Microsoft 365, Redmond, WA).

### Primary dermal fibroblast and keratinocyte culture

Dermal fibroblast and keratinocytes were isolated from P4 dorsal skin and activated/differentiated as previously described (Churko et al., 2011a, Churko et al., 2012, Churko et al., 2011b), but keratinocytes were cultured in KGM™ Gold Keratinocyte Growth Medium BulletKit™ (Lonza, Walkersville, MD). KO keratinocytes were grown on plates coated with 50ug/mL or 100ug/mL collagen, poly-D-lysine/laminin or human fibronectin (Corning, Bedford, MA). Brightfield images were taken on a ZEISS Axio Vert.A1 (ZEISS, Oberkochen, Germany). Live and dead cell percentages were determined using the Countess™ II automated cell counter (Thermo Fisher Scientific).

### Dermatitis incidence

Reports of dermatitis diagnosed by ACVS staff were counted for each genotype in this blinded, retrospective cohort study (Hampton et al., 2012). Mice were imaged using an iPhone XR (Apple, Inc., Cupertino, CA).

### Statistical analysis

Statistical analyses were performed using GraphPad Prism 9 (GraphPad Software, San Diego, CA).

## Supporting information

suppFigLeg

SuppMethods

## DATA AVAILABILITY STATEMENT

All raw data is available upon request.

## ACKNOWLEDGEMENTS

We would like to thank Dr. John Di Guglielmo (University of Western Ontario) for the gift of the TGF-β and Hisham Kamoun (University of Western Ontario) for his help with the Masson’s trichrome staining. We also thank Western University’s Animal Care and Veterinary Services staff for their care of the mice and providing the representative dermatitis image. Funding for this study was provided by the Natural Sciences and Engineering Research Council (NSERC) of Canada with a Discovery Grant (RGPIN-2015-06794) awarded to S. Penuela.

## CRediT Statement (author contributions)

Conceptualization: BO’D, SS, LD, SP; Data Curation: MK; Formal Analysis: BO’D, SS, MK, MB, CZ; Funding Acquisition: SP; Investigation: BO’D, RS-P, SS, CZ, DJ, JK, CBW; Methodology: KB; Project Administration: BO’D, SS, SP; Resources: KB, SP; Software: MK; Supervision: LD, SP; Visualization: BO’D, SS; Writing – Original Draft Preparation: BO’D, SP; Writing – Review and Editing: BO’D, RS-P, SS, MK, JK, LD, SS

---

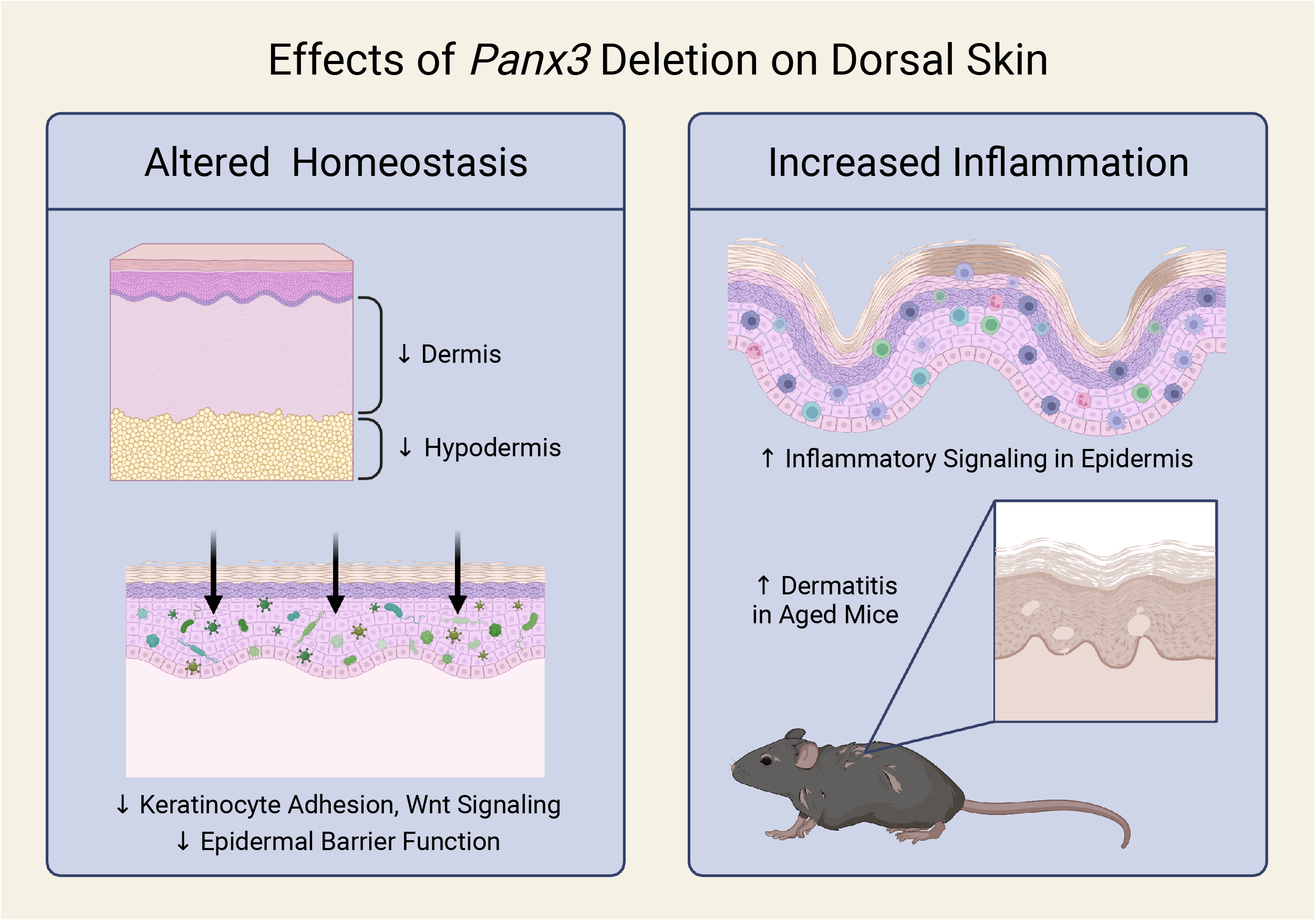

