## Supplementary figures and images for "Pannexin 3 channels regulate architecture, adhesion, barrier function and inflammation in the skin"

### FigS1

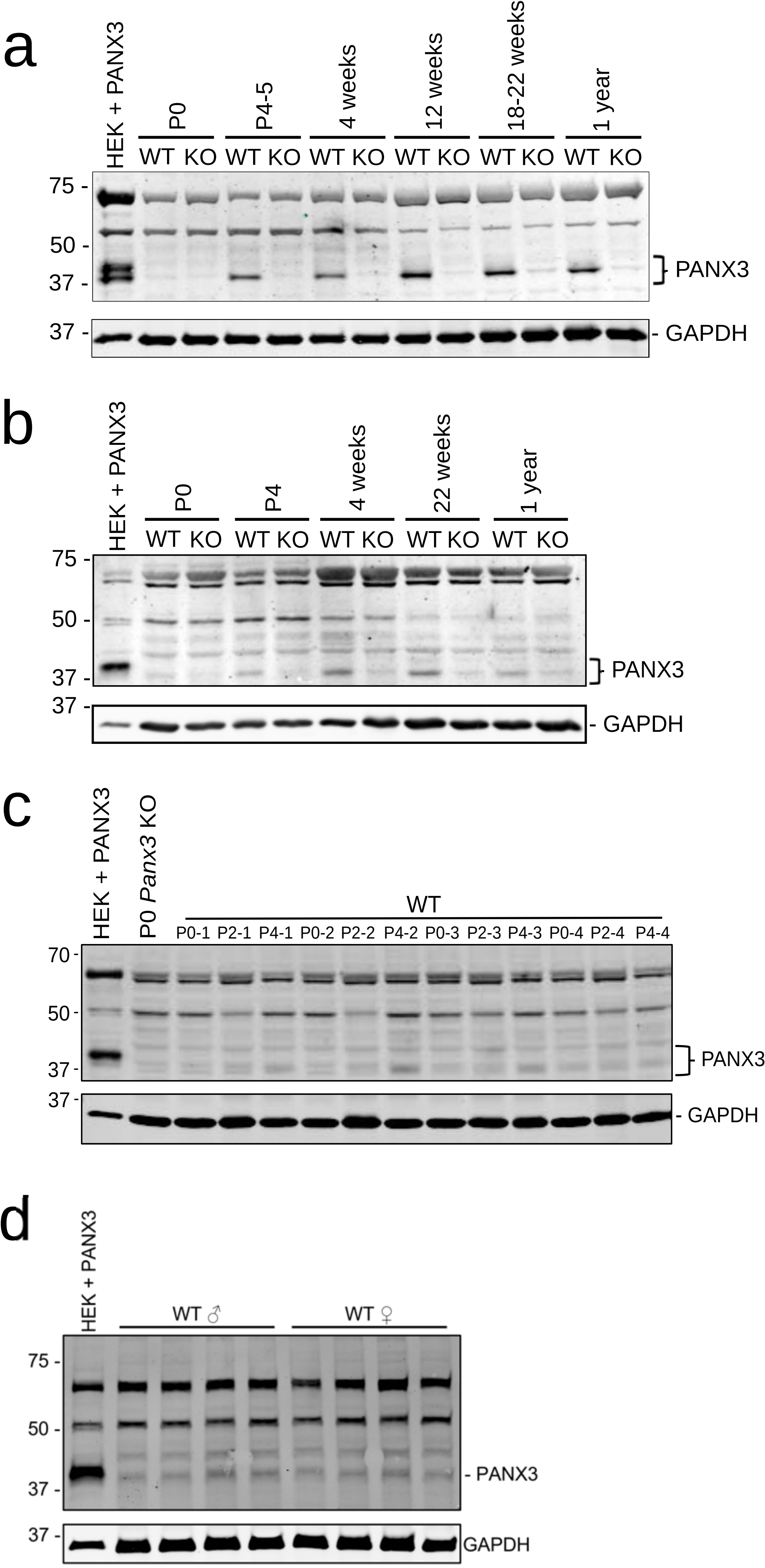

### FigS2

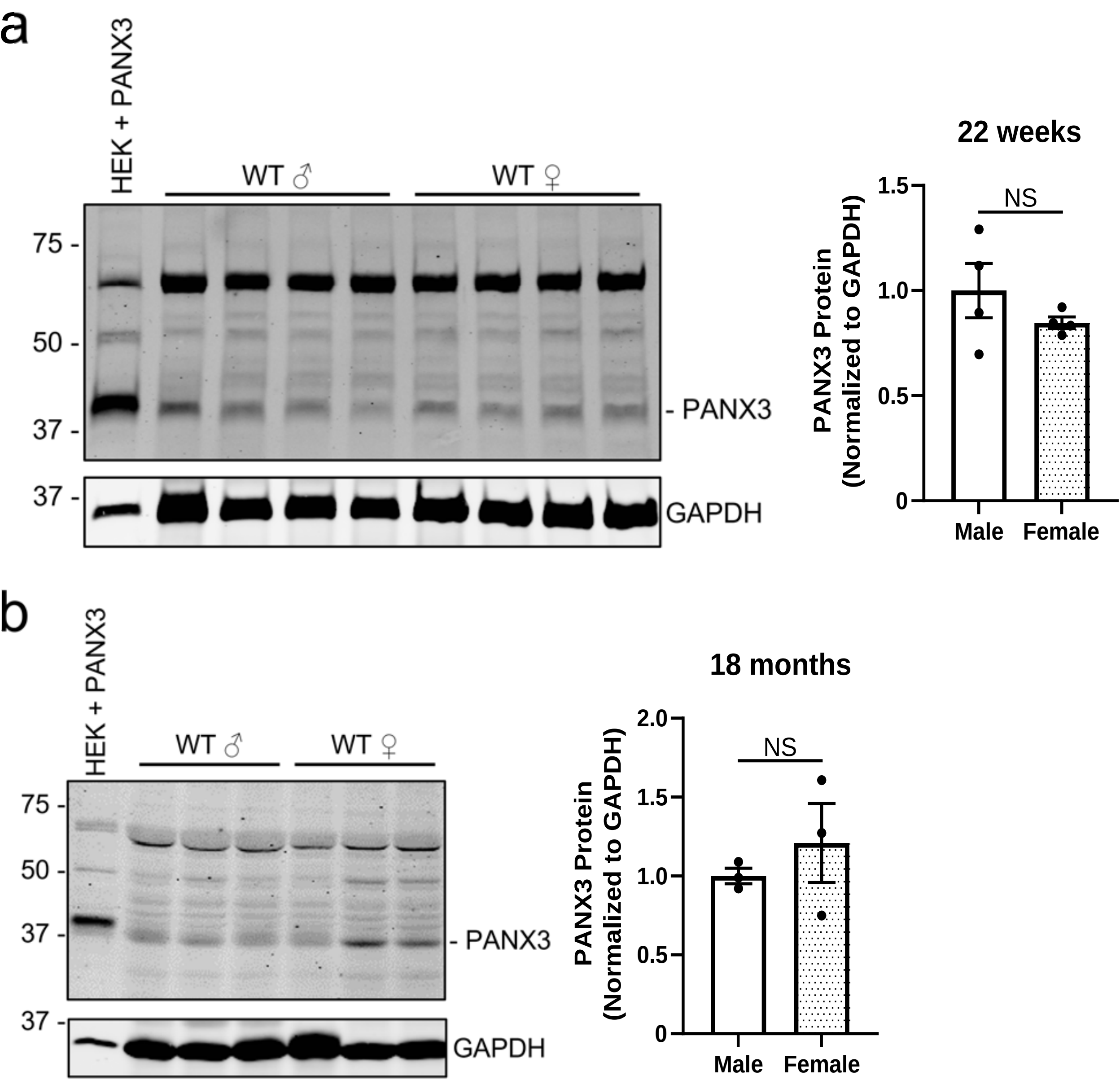

### FigS3

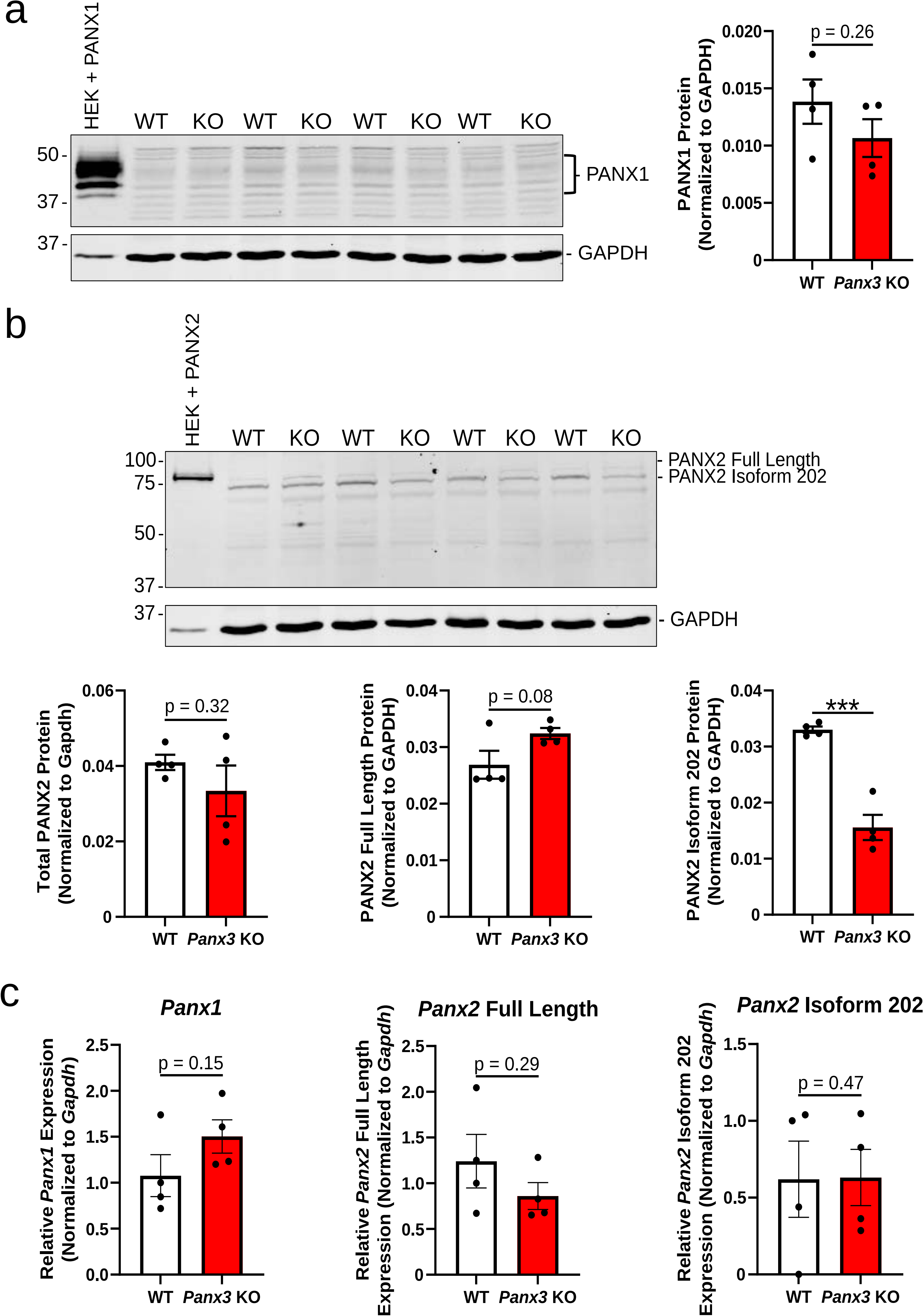

### FigS4

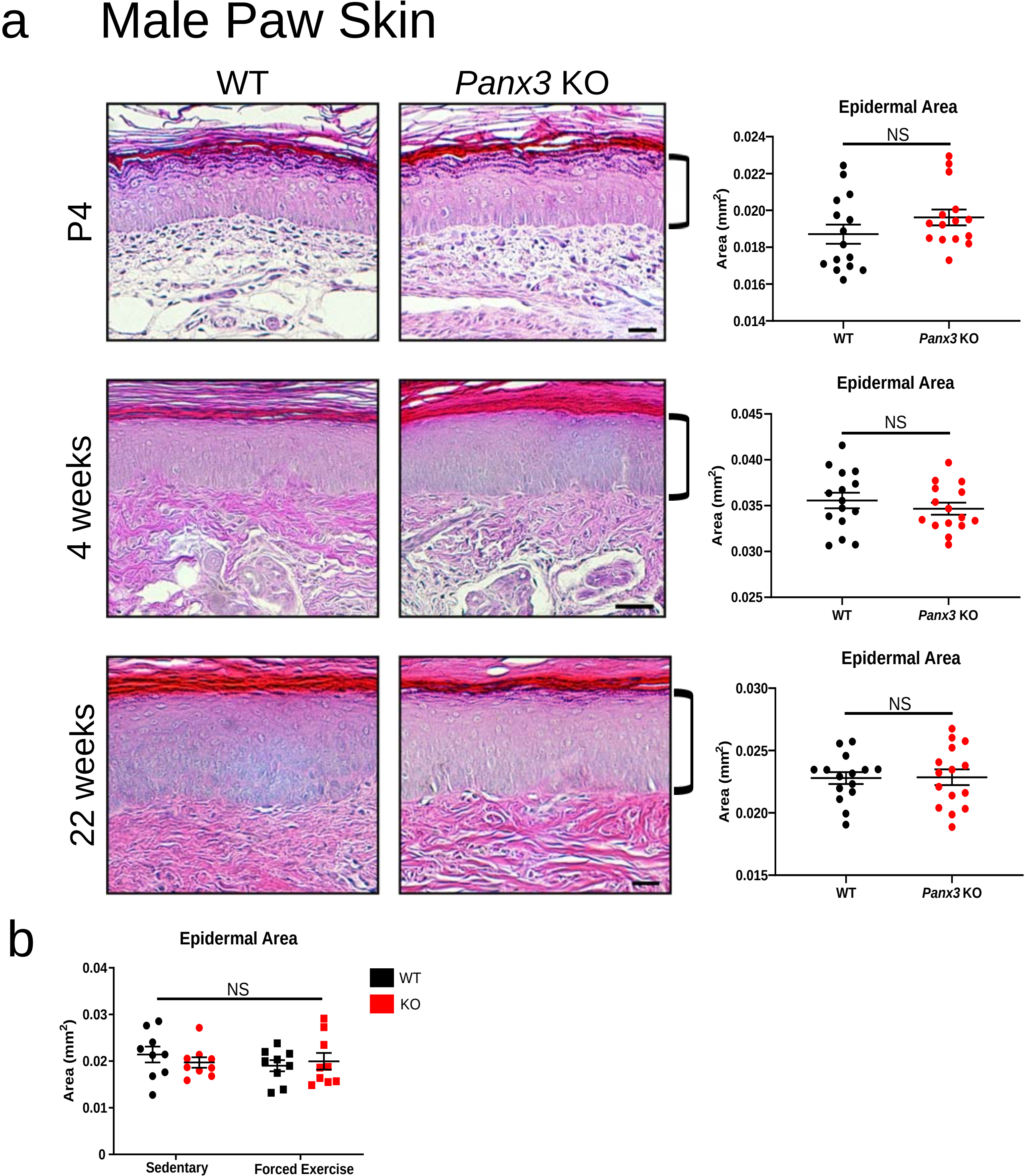

### FigS5

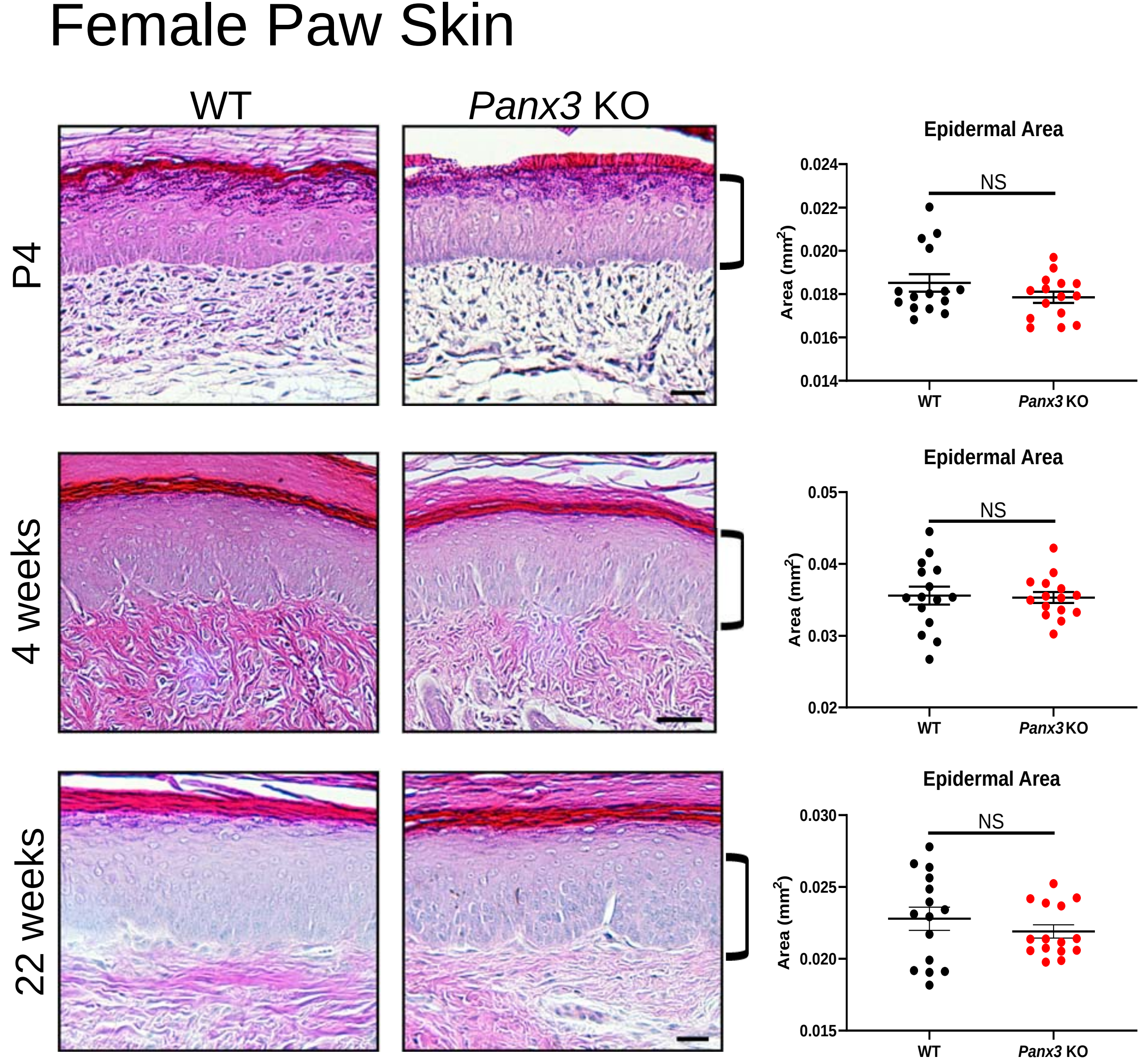

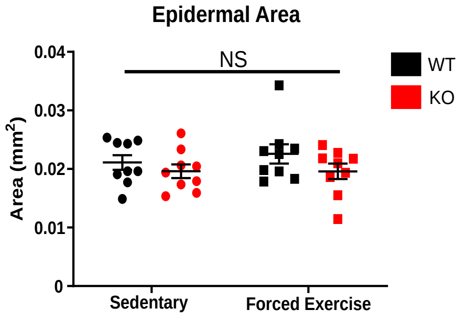

### FigS6

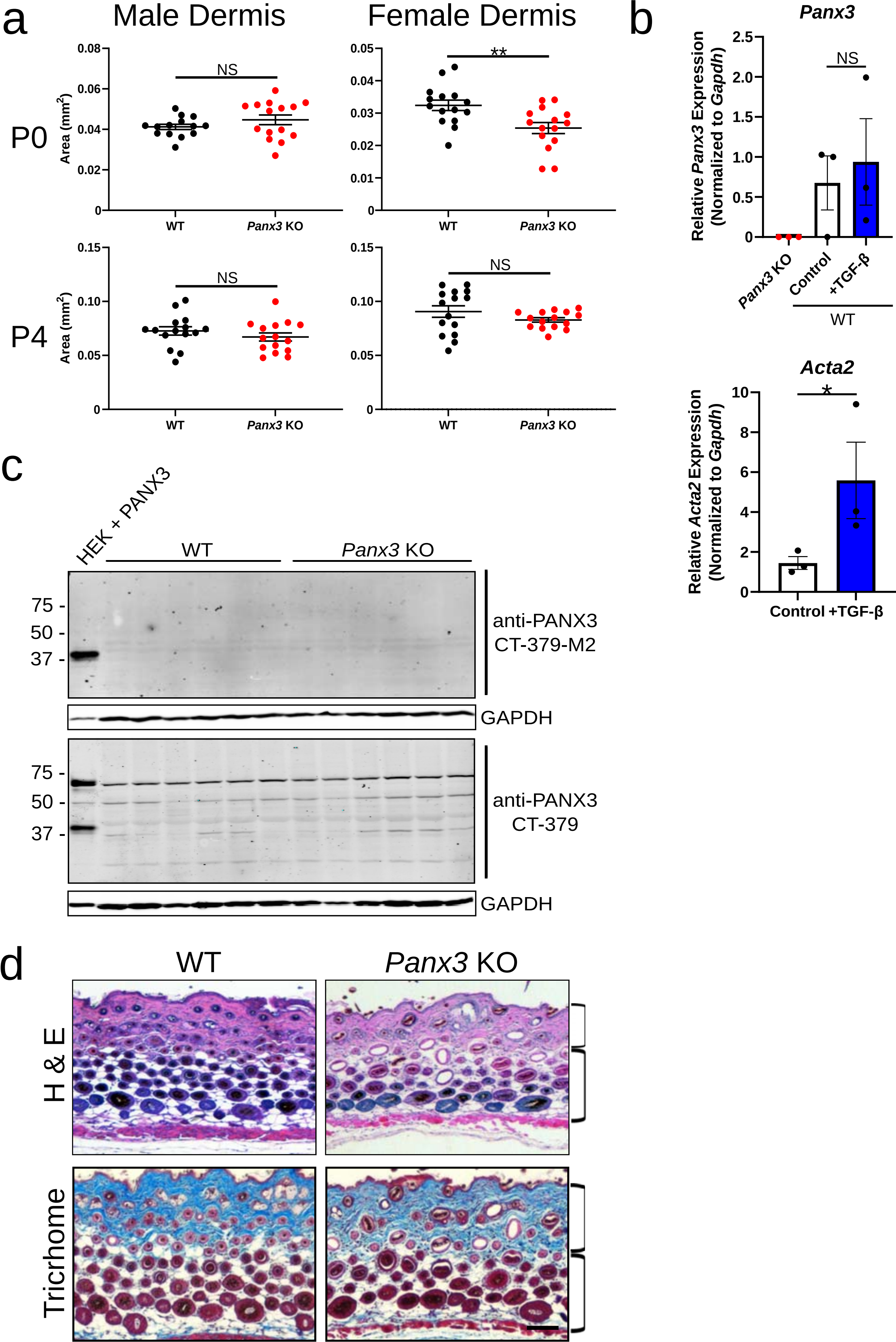

### FigS7

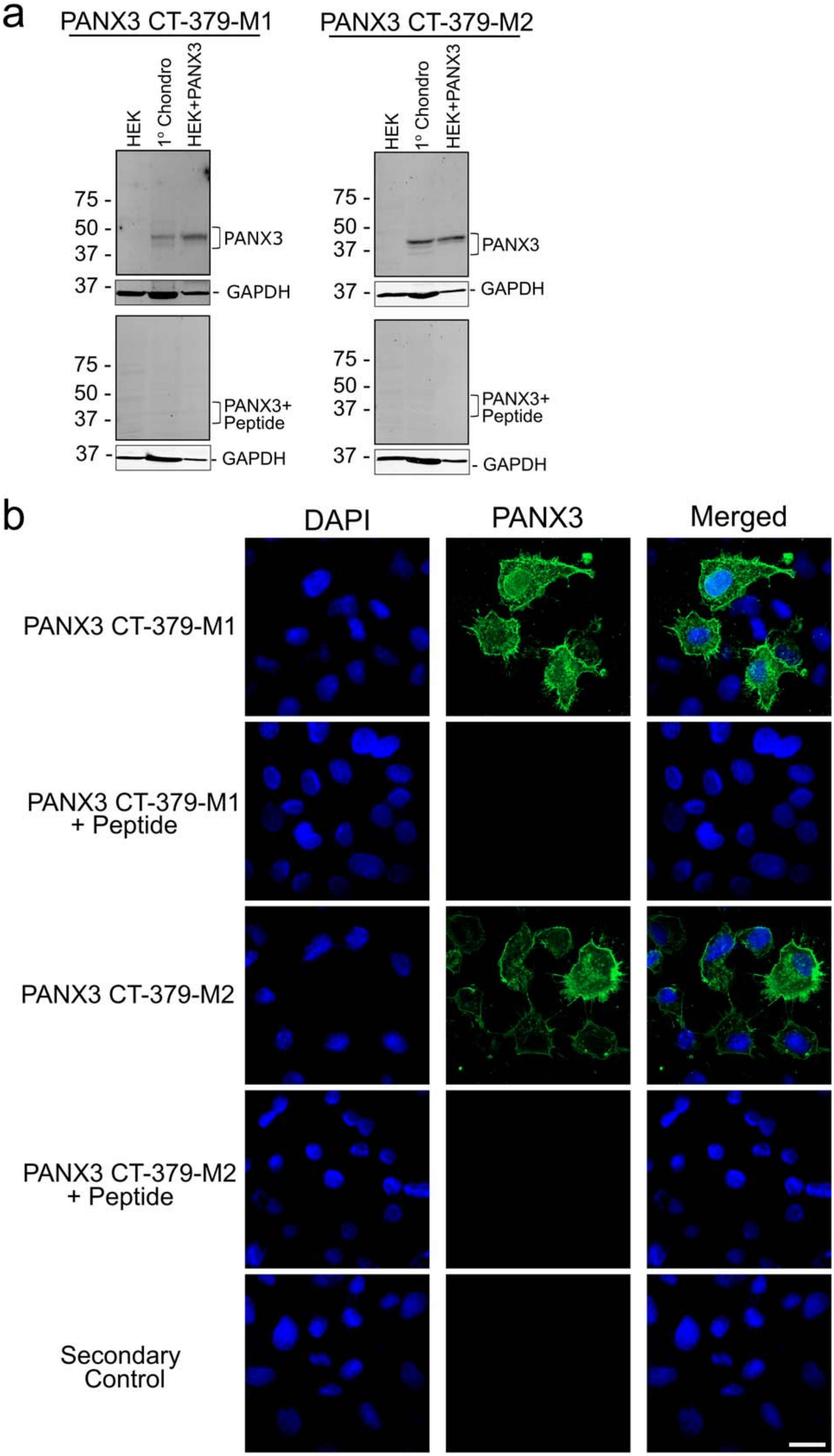

### FigS9

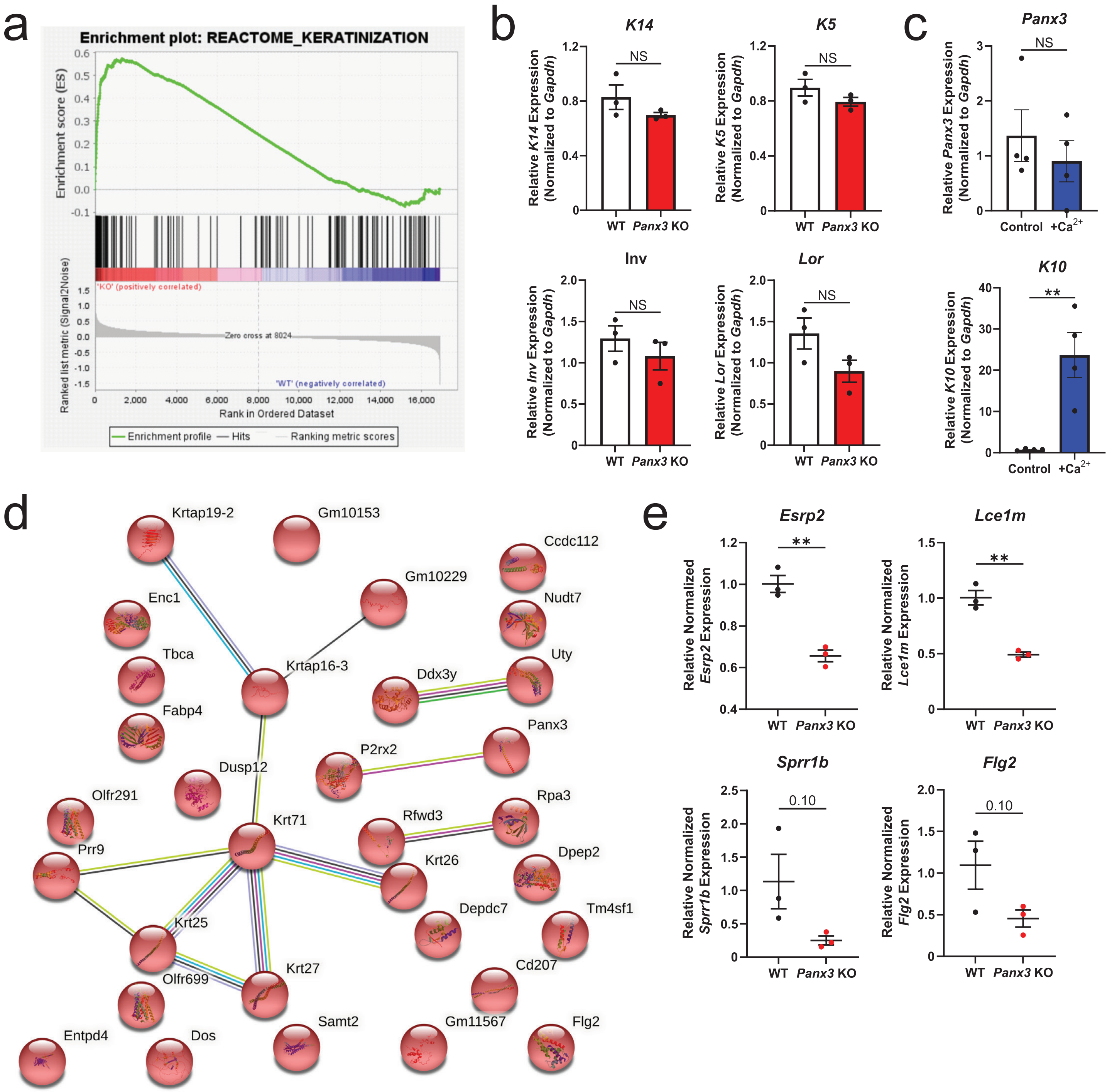
