## Supplementary material for "Pannexin 3 channels regulate architecture, adhesion, barrier function and inflammation in the skin": FigS8

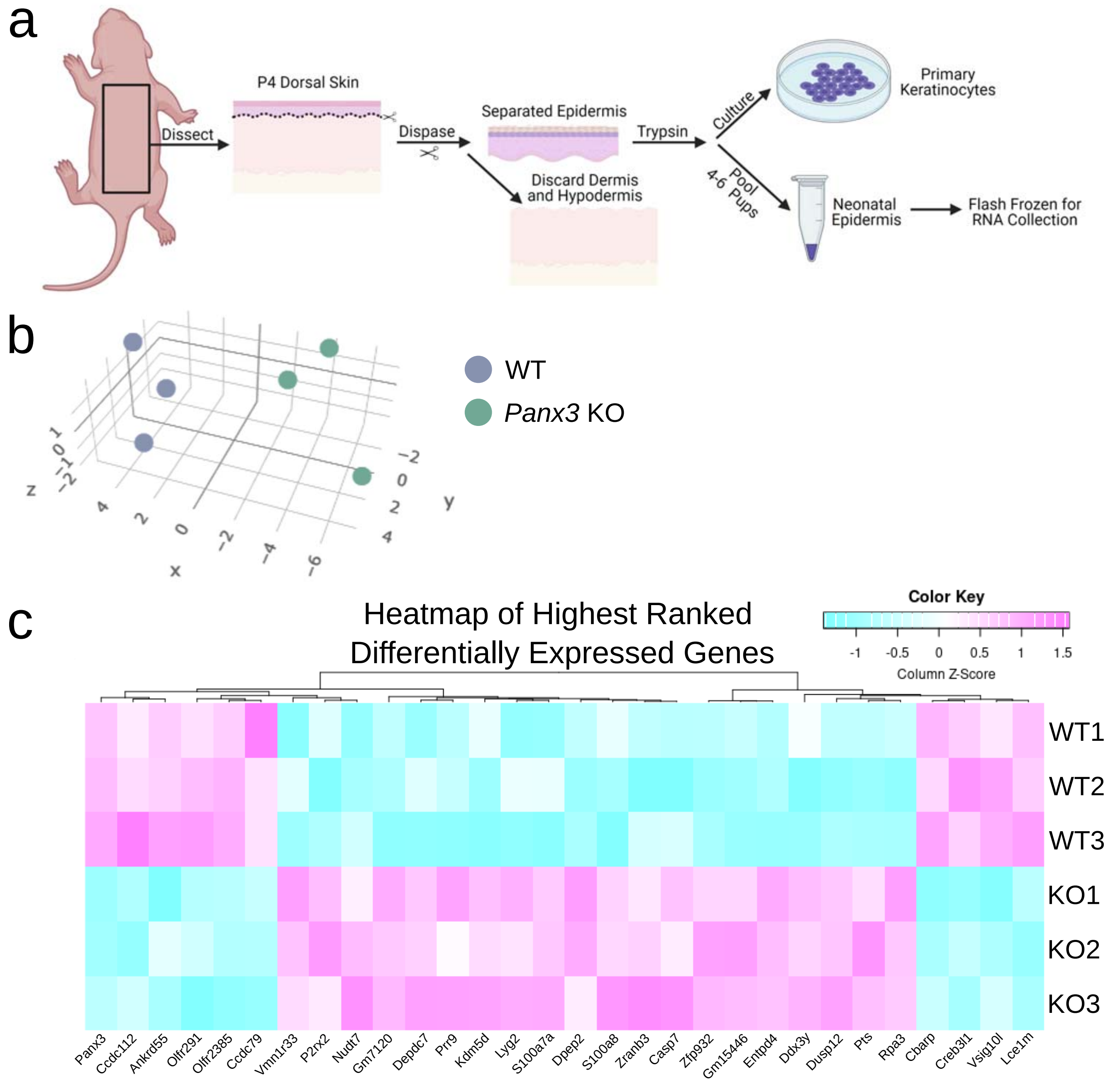


d

| Functional Category/Signaling Pathway | Gene Candidates with Altered Expression |
|---|---|
| Skin Development | *Stmn1, Casp7, Rptn, Krtap5-4, Entpd4* |
| Hair Follicles | *Stmn1, Lyg2, Casp7, Wfdc18, Fgfr1, Krtap4-8, Krt71, Tchh* |
| Wound Healing | *Gprc5a, Pts, Dpep2, Itm2a* |
| Barrier Function | *Esrp2, Lce1m, Stmn1, Sprr1b, Dgat2, Flg2, Elovl1* |
| Differentiation | *Arhgap32, Sh3kbp1, Cops3, Lce1m, Cpsf41, B9d1, Pts, Cyp2u1, Dynlt1, Rab4a, S100a7a, Sprr1b, H2afv, Krtap5-4* |
| Proliferation | *Rnf7, Rpa3, Pts, Cyp2u1, Stmn1, Lyg2, Pisd-ps3, Tchhl1, Tbca* |
| Cell-Cell and Cell-Matrix Adhesion | *Sh3kbp1, Cops3, Lrrc4c, Zmat3, Rab4a, Pard6a, Spink5, P3h2, Col6a4* |
| Inflammation, Psoriasis, Atopic Dermatitis and Infection | *Rgs1, Tsc22d3, Trat1, Il1rap, Dpep2, Il1r2, Pfdn4, Il18r1, S100a7a, Dpep2, P2rx2, S100a8, Casp7, Zranb3, Ccdc112, Cd207, Gm4553, Ltb4r2, Fabp4, Nlrp4e, Gas7, Crnn, Itm2a, Hpse, Atf3, Klk13, Gm19402, Lyz2, Irf1, Cbarp, Nudt7, Ankrd55, Lyg2* |
| Migration and Invasion | *Dynlt1, Gprc5a, Stmn1, Il1r2, Rpa3, Tm4sf1* |
| Metabolism | *Polr2h, Cox7a2l* |
| SCC | *Polr2h, Sh3kbp1, S100a7a, Fgfr1, Tchhl1* |
| Hedgehog Signaling | *Zfp932, B9d1, Pard6a* |
| Wnt/E-cadherin Signaling | *Il1r2, Pard6a, Arhgap32, Gpr137* |
| Smad Signaling | *Il1r2, Cdc42ep3* |
