## Supplementary material for "Pannexin 3 channels regulate architecture, adhesion, barrier function and inflammation in the skin": suppFigLeg

### SUPPLEMENTARY FIGURE LEGENDS

**Supplementary Figure S1. Full PANX3 blots.** Immunoblotting shows the presence of PANX3 in both male (**a**) and female (**b**) WT mouse dorsal skin from P4 to 1 year of age. (**c**) In female WT mice, PANX3 levels gradually increase from P0 to P4. (**d**) Immunoblotting shows no differences in PANX3 levels of WT male and female P4 dorsal skin. PANX3 levels normalized to GAPDH (protein loading control). All blots were probed with anti-PANX3 CT-379. In all samples, 50 kDa and 70 kDa immunoreactive species are present. Sizes in kDa.

**Supplementary Figure S2. PANX3 protein levels are consistent in 22 week and 1 year WT dorsal skin of both sexes.** Immunoblotting shows no differences in PANX3 levels of WT male and female 22 week (**a**) and 18 month (**b**) dorsal skin (NS, $p>0.05$; 22 week $N=4$, 18 month $N=3$). HEK cells ectopically expressing a mouse PANX3 plasmid (HEK + PANX3) were used as a positive control. PANX3 levels normalized to GAPDH (protein loading control). All blots were probed with anti-PANX3 CT-379. In all samples, 50 kDa and 70 kDa immunoreactive species are present. Sizes in kDa. Bars represent ±SEM. $N=4$. Unpaired t-test.

**Supplementary Figure S3. PANX2 isoform 202 levels are significantly decreased in KO dorsal skin.** Western blots with their respective quantifications show PANX1 (**a**) and PANX2 (**b**) levels in P4 WT and KO dorsal skin. There were no differences seen in PANX1, total PANX2 or full length PANX2 levels ($p>0.05$), but PANX2 isoform 202 was downregulated in KO dorsal skin (***$p<0.001$). HEK cells transfected with mouse pannexin plasmids were used as positive controls for PANX1 and full length PANX2 (HEK+PANX1, HEK+PANX2). GAPDH as protein loading control. Protein sizes in kDa. (**c**) mRNA levels of *Panx1* and *Panx2* were not significantly different between the dorsal skin of each genotype ($p>0.05$). *Gapdh* was used to calculate normalized mRNA expression via ΔΔCT. Bars represent ±SEM. $N=4$. Unpaired t-test.

**Supplementary Figure S4. Male WT and KO thick skin epidermal area are not significantly different in sedentary and forced exercise conditions.** (**a**) Histological analysis and epidermal area (minus the corneal layer) measurements were performed on male murine paw skin at P4, 4 weeks and 22 weeks of age. No significant differences were seen between the epidermal areas of male WT and KO mice at any timepoint ($p>0.05$, unpaired t-test). $N=5$, $n=15$. Scale bars: 25µm. (**b**) The epidermal area of male WT and KO paws subjected to forced exercise did not differ from each other or those of sedentary mice ($p>0.05$, two-way ordinary measures ANOVA). $N=3$, $n=9$. Error bars represent ±SEM. NS, no significance.

**Supplementary Figure S5. The paw skin epidermal area of female WT and KO mice show no significant differences, even after forced exercise.** (**a**) Histological analysis and epidermal area (minus the corneal layer) measurements were performed on female murine paw skin at P4, 4 weeks and 22 weeks of age. Female KO paw epidermal area did not differ from that of female WT paws at any age examined ($p>0.05$, unpaired t-test). $N=5$, $n=15$. Scale bars: 25µm. (**b**) The epidermal area of female WT and KO paws subjected to forced exercise did not differ from each other or those of sedentary mice ($p>0.05$, two-way ordinary measures ANOVA). $N=3$, $n=9$. Error bars represent ±SEM. NS, no significance.

**Supplementary Figure S6. PANX3 is undetectable in P4 WT dermal fibroblasts.** (**a**) Histological analysis revealed P0 female dorsal skin dermal area was significantly reduced compared to controls (**$p<0.01$, N=5, n=15), but unchanged in all other measurements (unpaired t-test). (**b**) PANX3 protein was undetectable via immunoblotting in dermal fibroblasts isolated from P4 WT and KO dorsal skin ($N=6$). GAPDH as protein loading control, HEK+PANX3 as positive control, protein sizes in kDa. (**c**) No differences in (mRNA) *Panx3* in WT fibroblasts in control or TGF-β conditions via RT-qPCR ($N=3$, $p>0.05$; one-way ordinary measures ANOVA).

Fibroblast activation was confirmed by α-smooth muscle actin (*Acta2*) upregulation (*$p<0.05$, paired t-test). *Gapdh* was used to calculate normalized mRNA expression via ΔΔCT. Bars represent ±SEM. (**d**) Masson's Trichrome staining of 4-week-old male WT and KO mouse dorsal skin show similar levels of collagen abundance. Histological staining (H&E) also shown for reference. *N*=3, scale bar: 100μm. NS, no significance.

**Supplementary Figure S7. Characterization of PANX3 CT-379-M1 and M2 antibodies.** (**a**) Western blotting using non-transfected HEK cells, primary mouse chondrocytes (1° chondro) and HEK cells ectopically expressing mouse PANX3 (HEK+PANX3). Blots were probed with 1:100 PANX3 CT-379-M1 or PANX3 CT-379-M2, peptide pre-adsorption assay confirmed antibody specificity. GAPDH as a loading control. Protein sizes in kDa. (**b**) Confocal micrographs of NRK cells overexpressing mouse PANX3. Cells were immunolabeled with 1:50 dilutions of PANX3 CT-379-M1 or PANX3 CT-379-M2 (green) with or without peptide pre-adsorption. Nuclei (blue) were counterstained with Hoescht 33342. Scale bar: 20μm. The peptide was pre-incubated with each antibody for 30 minutes, in 50:1 molar excess of each antibody, and ran in parallel to the antibody minus peptide conditions.

**Supplementary Figure S8. Analysis of WT and *Panx3* KO neonatal epidermis Clariom™ S expression profiling data.** (**a**) Schematic (Biorender.com) showing primary keratinocytes and neonatal epidermis isolation from P4 pups (**b**) Clariom™ S transcriptomic analysis of WT and KO P4 neonatal epidermis (*N*=3, pooled from 4-6 pups, mix of male and female) was analyzed by RStudio Cloud. A principal component analysis visualized expression differences in the 50 highest ranked genes shows sample separation by genotype. (**c**) Heat map showing the 30 highest ranked genes with fold change of minimum ±2 column Z-score and $p<0.05$ between WT and KO neonatal epidermis. (**d**) Major functional categories and signaling pathways affected by *Panx3* ablation in neonatal epidermis and the corresponding gene candidates identified to have altered expression in KO samples.

**Supplementary Figure S9. Differentially expressed genes in KO neonatal epidermis function in epidermal barrier formation and maintenance.** (**a**) Keratinization was the most positively enriched gene set in KO neonatal epidermis (Enrichment Score, ES>0) via GSEA pathway analysis (*N*=3). (**b**) No differences were seen in undifferentiated (*K14, K5*) or differentiated (*Inv, Lor*) keratinocyte marker transcripts between neonatal epidermis between genotypes (NS, no significance, *N*=3, unpaired t-test). (**c**) No differences were seen in *Panx3* transcript levels in WT keratinocytes cultured in control or $Ca^{2+}$ differentiation conditions (*N*=4, NS, paired t-test). Keratinocytes differentiation was confirmed by an increase in *K10* transcript (*p*=0.0012). *Gapdh* was used to calculate normalized mRNA expression via ΔΔCT. (**d**) STRING plots were generated using the first 100 highest ranked differentially expressed genes showing ±2-fold expression changes. This cluster was associated with keratinization, epidermis morphogenesis and formation of the cornified envelope. (**e**) Epithelial splicing regulatory protein 2 (*Esrp2*) and late cornified envelope 1m (*Lce1m*) transcript levels were significantly reduced and cornifin-B (*Sprr1b*) and filaggrin 2 (*Flg2*) trended to a reduction in KO neonatal epidermis compared to WT (**$p<0.01$, unpaired t-test). All are gene candidates associated with epidermal barrier function. Expression normalized and presented relative to the WT average (*N*=3). Bars represent ±SEM.
