## Supplementary material for "Pannexin 3 channels regulate architecture, adhesion, barrier function and inflammation in the skin": SuppMethods

### SUPPLEMENTAL MATERIALS AND METHODS

#### Animals and animal ethics

Global *Panx3* KO mice used were previously generated by our group and congenic C57BL/6N (WT) mice were used as controls (Moon et al., 2015). Mice were genotyped to verify the presence or lack of exon 2 in *Panx3* which requires two separate PCR reactions (see Table S1 for primers). After mouse euthanasia, tail tips were digested in a 0.2 mM EDTA, 25 mM NaOH solution pH 8 and then the solution was neutralized using a 40 mM Tris-HCl pH 5 solution. The Platinum® *Taq* DNA High Fidelity Polymerase (11304011; Invitrogen, Carlsbad, CA) and manufacturer's protocol was used for both reactions, with an annealing temperature and extension time of 58°C for 1.15 min for reaction 1 (R1) and 56°C for 1 min for reaction 2 (RT). PCR products were separated on a 1% agarose gel. R1 is used to distinguish between WTs and mice with at least one mutant allele, yielding a 1326 bp WT DNA fragment or a 600 bp DNA fragment in heterozygotes or KOs. RT distinguishes a heterozygous mouse from a full KO, where WT and heterozygous mice produce a 770 bp band, absent in a *Panx3* KO mouse. All animal experiments followed guidelines and protocols approved by the Animal Care Committee at the University of Western Ontario (#2019-069). Mice were fed Teklad 2018 (Envigo, Indianapolis, IN) *ad libitum* and maintained in a 12-hr-light to 12-hr-dark cycle.

#### Protein extraction and western blotting

Dorsal skins were dissected from WT and *Panx3* KO mice at P0, P2, P4, 4 weeks, 12 weeks, 22 weeks, 1 year and 18 months, pulverized in liquid nitrogen and protein was extracted using a 1x radioimmunoprecipitation assay buffer (50mM Tris-HCl pH 8.0, 150 mM NaCl, 1% Nonidet® 40 Substitute (74385; Sigma-Aldrich, St. Louis, MO), 0.5% sodium deoxycholate) containing 1 mM

NaF, 1mM $Na_3VO_4 \cdot 2H_2O$ and a quarter of a Pierce™ Protease Inhibitor EDTA-free Mini Tablets (A32955; Thermo Fisher Scientific, Rockford, IL), as previously described (Abitbol et al., 2019). Protein concentrations were determined using the Pierce™ BCA Protein Assay Kit (23225; Thermo Fisher Scientific, Rockford, IL). After sample protein separation using 10% SDS-PAGE gels (40 µg per sample for all blots), membranes were transferred to a nitrocellulose membrane via the iBlot™ Gel Transfer Device (Invitrogen, Carlsbad, CA). Membranes were blocked overnight with 3% BSA (ALB001.100; BioShop, Burlington, Canada) in 1x PBS. Blots were incubated with primary antibodies diluted in 1x PBS with 0.05% Tween20 overnight at 4°C or 1 hr at room temperature for GAPDH. Primary antibody concentrations included: anti-PANX1 CT-395 0.5 µg/mL, anti-PANX2 CT-523 4 µg/mL and anti-PANX3 CT-379 1 µg/mL (Penuela et al., 2009); generated anti-PANX3 CT-379-M1 and M2 unpurified supernatant 1:100, and anti-mouse GAPDH 0.2 µg/mL (Millipore Sigma, Burlington, MA, G8795; protein loading control). IRDye®-800CW goat anti-rabbit and -680RD goat anti-mouse IgG secondary antibodies at a 0.1 µg/mL concentration (LI-COR Biosciences, Lincoln, NE, 926-32211 and 926-68070 respectively) were used for signal detection and blots were imaged on a LI-COR Odyssey Infrared Imaging System (LI-COR Biosciences, Lincoln, NE). Protein levels were quantified using ImageStudio (LI-COR Biosciences, Lincoln, NE), normalized to GAPDH, and presented relative to one WT mean value. For PANX1, PANX2 and PANX3 positive controls, human embryonic kidney 293T cells (HEK; CRL-3216; ATCC, Manassas, VA; RRID: CVCL_0063) were cultured in DMEM media (12430062; Gibco™, Grand Island, NY) with 10% FBS (098-150; WISENT Inc., Saint-Jean-Baptiste, Canada), 1% penicillin/streptomycin (15140-122; Gibco™, Grand Island, NY) and transfected with 5.5 µg mouse PANX1, PANX2 or PANX3 plasmids in a 10 cm plate (Penuela et al., 2007, Penuela et al., 2009) using Lipofectamine 3000 (Invitrogen, Carlsbad, CA) according to the manufacturer's protocol.

#### Histology and Masson's trichrome staining

Histological staining and measurements were performed on both male and female, WT and *Panx3* KO dorsal skin and paw skin of various ages as described previously (Abitbol et al., 2019). Briefly, tissue was fixed overnight at 4°C in 10% neutral-buffered formalin, processed at the Robart's Research Institute Molecular Pathology Core Facility (University of Western Ontario), embedded in paraffin and sectioned. Parallel sections (5-7 µm) were deparaffinized, rehydrated and stained with Mayer's hematoxylin (MHS16; Sigma-Aldrich, St. Louis, MO) and eosin-Y (1931492; Lerner Laboratories, Pittsburgh, PA). Stained slides were mounted using Fisher Chemical™ Permount™ Mounting Medium (SP15; Fisher Scientific, Pittsburgh, PA) and imaged. ImageJ (Fiji) (Schindelin et al., 2012) was used to crop each image to a standard frame area, manually segment and then measure the epidermal and dermal area (minus the *stratum corneum*) and hypodermal areas of the dorsal skin and the epidermal area (minus the corneal layer) of the paw skin. Four-week-old male dorsal skin sections were stained with Masson's trichrome stain following a previously reported protocol (Robb et al., 2021). Briefly, paraffin sections were deparaffinized with xylene, rehydrated in a descending ethanol series and then stained with Bouin's solution (HT10132; Sigma, St. Louis, MO), Weigert's iron hematoxylin A and B (26044-05 and 26044-15; Electron Microscopy Sciences, Hatfield, PA), Biebrich Scarlet-Acid Fucshin (HT151; Sigma, St. Louis, MO), 10% phosphotungstic acid/10% phosphomolybdic acid (19500/19400; Electron Microscopy Sciences, Hatfield, PA) and then Aniline Blue (B8563; Sigma, St. Louis, MO) before incubation with 1% glacial acetic acid, ethanol dehydration and mounting (as above). Hematoxylin and eosin (H&E) and Masson's trichrome stained slides were imaged on a Leica DMIL LED brightfield inverted microscope (Leica, Wetzlar, Germany) at either 5x or 20x magnification.

#### Forced exercise protocol

The forced exercise protocol used is outlined previously (Wakefield et al., 2021). Briefly, 24-week-old male and female WT and *Panx3* KO mice were randomized to sedentary or forced exercise groups. Mice in the forced exercise group were forced to run on a treadmill (Columbus Instruments, Columbus, OH) for 1 h daily, 5 days per week in a 6-week period at a speed of 11 m/s and 10º incline. Then hind paws were dissected from sedentary and forced exercised mice, fixed, stained, and analyzed as above, except the paw epidermal areas were manually segmented and measured by a blind observer.

#### RNA isolation and RT-qPCR

Following mouse euthanasia, P4 and 22 week dorsal skin was dissected from WT and *Panx3* KO mice and immediately flash-frozen in liquid nitrogen. Ribonucleic acid (RNA) was extracted from dorsal skin using a hybrid TRIzol™ (15596018; Life Technologies, Carlsbad, CA) and RNeasy Plus Mini Kit (74134; Qiagen, Venlo, Netherlands) protocol as previously described (Abitbol et al., 2019). For primary cells and neonatal epidermis, the RNeasy Plus Mini Kit (74134; Qiagen, Venlo, Netherlands) was used for RNA extraction. Fifty ng of RNA was then reverse transcribed into cDNA using a High-Capacity cDNA Reverse Transcription Kit with RNase Inhibitor (4374966; Thermo Fisher Scientific, Rockford, IL) and following the manufacturer's instructions. Reverse transcriptase-quantitative polymerase chain reaction (RT-qPCR) was performed using

SsoAdvanced Universal SYBR® Green Supermix (1725274; Bio-Rad, Hercules, CA) according to the manufacturer's protocol. See Table S1 for a list of primers used. Samples were run in duplicate. Transcript abundance was normalized to *Gapdh*, presented as relative to one WT mean value and analyzed using the ΔΔCT method in Excel (Microsoft 365, Redmond, WA). While geometric means were represented in the graphs, logarithmic means were used to perform statistical analyses.

**Generation and characterization of PANX3-specific antibodies**

Carboxyl-terminal amino acid residues 379-392 (KPKHLTQHTYDEHA) of mouse PANX3 were used in mouse monoclonal antibody generation by Genemed Biotechnologies, Inc. (Torrance, CA). Hybridoma supernatant from ten generated clones were tested via immunoblotting, and two clones with the highest specificity and signal intensity were selected and designated PANX3 CT-379-M1 and PANX CT-379-M2. The unpurified supernatant of PANX3 CT-379-M2 was used in the dermal fibroblast Western blot analysis alongside the previously published and characterized PANX3 CT-379 (Penuela et al., 2007). Characterization of these monoclonal antibodies by immunofluorescence (1:50 dilution) and immunoblotting (1:100 dilution) with peptide pre-adsorption as described previously (Freeman et al., 2019) is shown in Figure S6. The peptide was pre-incubated with each antibody for 30 minutes, in 50:1 molar excess of each antibody, and ran in parallel to the antibody minus peptide conditions. For immunoblotting, IRDye®-680RD goat anti-mouse IgG secondary antibody at a 0.1 µg/mL concentration was used. Primary mouse chondrocytes were obtained from WT mice as described previously (Gosset et al., 2008), and protein was isolated as described above. For immunocytochemistry, Normal Rat Kidney (NRK; CRL-6509; ATCC, Manassas, VA).; RRID: CVCL_3758) cells were plated on coverslips in a 6 well plate and transfected with 1 µg of mouse PANX3 plasmid using Lipofectamine 3000 according to the manufacturer's protocol. Forty-eight hours after transfection, cells were washed with 1x DPBS without $Ca^{2+}$ or $Mg^{2+}$ (14190250; Gibco™, Grand Island, NY) and fixed with ice-cold methanol:acetone (5:1, vol/vol) for 15 min at 4°C. Coverslips were blocked for 1 hr at room temperature with 10% goat serum (50062Z; Life Technologies, Carlsbad, CA), incubated with the primary monoclonal antibodies diluted in 1% goat serum overnight at 4°C, washed with DPBS without $Ca^{2+}$ or $Mg^{2+}$ and then incubated with Alexa Fluor™ 488 goat anti-mouse secondary antibody (1:700; A11029; Invitrogen, Carlsbad, CA) for 1 hr at room temperature. After washing, Hoechst 33342 (H3570; Life Technologies, Carlsbad, CA) diluted 1:1000 in double-distilled water was incubated with the samples for 5 min at room temperature to stain cell nuclei and Aqua-Mount™ (13800; Lerner Laboratories, Pittsburgh, PA) was used to mount the coverslips. Immunofluorescence images were obtained using a ZEISS LSM 800 AiryScan Confocal Microscope from the Schulich Imaging Core Facility (University of Western Ontario) using 405 nm (Hoechst 33342) and 488 nm (Alexa Fluor 488) laser lines.

**Epidermal barrier function assay**

The epidermal barrier function of P2 WT and *Panx3* KO male and female pups was assessed using toluidine blue staining previously described (Press et al., 2017). Pups were euthanized using $CO_2$, incubated briefly in ascending and then descending concentrations of methanol solutions (25%, 50%, 75%, 100%; diluted in water) and stained with 0.2% toluidine blue (89640; Sigma-Aldrich, St. Louis, MO) for 15 min. After three rinses in 90% ethanol and a wash in water, pups were imaged using an iPhone SE (Apple, Inc., Cupertino, CA) and examined for blue dye permeation.

Intentional barrier disruption was performed using a scalpel to make a small incision in the dorsal skin.

**Clariom$^{TM}$ S expression profiling and analysis**

RNA from 3 pooled samples each (4-6 pups, includes mix of males and females) of P4 WT and *Panx3* KO neonatal epidermis were sent for Clariom$^{TM}$ S, Mouse expression profiling at the Genetic and Molecular Epidemiology Laboratory at McMaster University. Expression profiling data was analyzed using TAC version 4.0.2.15 (Thermo Fisher Scientific, Rockford, IL) and RStudio Cloud (https://www.rstudio.com/products/cloud/) version 4.1.2 (RStudio Team, 2022) software to find DEGs between WT and *Panx3* KO neonatal epidermis where differential expression was represented by a gene-level fold change of minimum ±1.5 and $p<0.05$ in *Panx3* KO samples compared to controls. GSEA was used to investigate pathway-level changes by comparing the gene lists produced by Clariom S$^{TM}$ expression profiling against *a priori* defined gene sets provided by GSEA and MSigDB C2 collection version 7.4 (Mootha et al., 2003, Subramanian et al., 2005). Enrichment Scores (ES) were generated for each gene set using a weighted Kolmogorov-Smirnov-like statistic (Subramanian et al., 2005). The ES demonstrates the degree that genes in a specific gene set are positively or negatively enriched in the *Panx3* KO neonatal epidermis compared with WT. STRING plots (https://string-db.org/) were generated using the first 100 highest ranked differentially expressed genes showing ±2-fold expression changes, setting the organism to *mus musculus* or projecting the *mus musculus* genes to *homo sapiens*, and kmeans clustering into three clusters (Szklarczyk et al., 2019, Szklarczyk et al., 2021). Reactome (https://reactome.org/) pathway analysis was also performed using minimum ±1.5-fold change gene sets with all mouse genes converted to their human-equivalent (Fabregat et al., 2016, Fabregat et al., 2017).**Primary dermal fibroblast and keratinocyte culture**

P4 dorsal skin was dissected from WT and *Panx3* KO male and female pups for primary dermal fibroblast and keratinocyte isolation. *Panx3* KO dermal fibroblasts were isolated using 5 cu/mL dispase (354235; Corning, Bedford, MA) and 2 mg/mL collagenase type I (LS004196; Worthington Biochemical Corporation, Lakewood, NJ) treatment, cultured in DMEM media with 10% FBS and 1% penicillin/streptomycin (as above), and activated into myofibroblasts by TGF-β stimulation as previously described (Churko et al., 2011b). Briefly, 1.5 x $10^5$ cells were seeded on collagen-coated 60 mm dishes and treated daily with 200 pM TGF-β1 (100-21; PeproTech, Cranbury, NJ) dissolved in DMEM media with 1% BSA and 1% penicillin/streptomycin for 4 days. Activation was confirmed by an increase in *Acta2* transcripts. Primary keratinocytes were isolated following the already established dispase and 0.25% trypsin/1 mM EDTA (325-043-EL; WISENT Inc., Saint-Jean-Baptiste, Canada) protocol (Churko et al., 2011a), but were cultured in KGM™ Gold Keratinocyte Growth Medium BulletKit™ (192060; Lonza, Walkersville, MD) on uncoated tissue culture plastic (see schematic in Fig. 7A). Trypsin was neutralized using an equal volume of 0.5 mg/mL soybean trypsin inhibitor (17075-029; Gibco™, Grand Island, NY). *Panx3* KO keratinocytes isolated from 3 male and 3 female pups (*N*=6) were also grown on tissue culture plates coated with 50 ug/mL or 100 ug/mL rat tail type I collagen (354236; Corning, Bedford, MA), poly-D-lysine/laminin (354595; Corning, Bedford, MA) or human fibronectin (354402; Corning, Bedford, MA). Brightfield images of primary keratinocytes were taken at 20x magnification using a ZEISS Axio Vert.A1. (ZEISS, Oberkochen, Germany). For morphological analysis, three separate isolations (from three different litters) per genotype were performed where keratinocytes isolated from 3 pups per sex were imaged (*N*=18 total, *N*=9 per sex). WT keratinocytes were differentiated with 1.4 mM $CaCl_2$ treatment, confirmed by an increase in *K10* transcripts (Churko et al., 2012). For neonatal epidermis samples, the Churko *et al*. (2011a) isolation protocol was followed, except at the last step where cells would normally be cultured, un-plated epidermis samples from 4-6 pups (includes mix of both sexes) were pooled, pelleted via centrifugation and flash frozen in liquid nitrogen for RNA isolation. Live and dead cell percentages of WT and *Panx3* KO un-plated epidermis samples were determined using the Countess™ II automated cell counter (Thermo Fisher Scientific, Rockford, IL) and trypan blue (ICN1691049; Fisher Scientific, Pittsburgh, PA). Measurements were performed on processed neonatal epidermis samples (*N*=9, cells isolated from 5 male and 4 female pups) immediately before the cells were plated to rule out differences in cell damage due to the isolation procedure.

**Dermatitis incidence**

To investigate the incidence of dermatitis in an aging mouse colony we analyzed reports of dermatitis cases diagnosed by Western University's ACVS staff using their SAR Treatment Algorithm for Mouse in a blinded, retrospective cohort study (Hampton et al., 2012). A total of 118 aged mice, constituting 62 WT (22 males and 40 females) and 56 *Panx3* KO (26 males and 30 females) mice, were included in this retrospective cohort study and ranged from 10.5-18 months of age at the time of diagnosis. The number of mice in each sex and genotype diagnosed with dermatitis by a blinded veterinary assessment were counted and used to calculate the incidence of dermatitis as well as the odd's ratio of developing dermatitis. Mice were imaged using an iPhone XR (Apple, Inc., Cupertino, CA), courtesy of ACVS.

**Statistical analysis**

Statistical analyses were performed using GraphPad Prism 9 (GraphPad Software, San Diego, CA) with specific test details in each figure legend. Error bars represent mean ± SEM. All data shown is representative of three to five independent experiments or mice with two to three technical replicates, except for Fisher's exact tests for dermatitis incidence. **REFERENCES**

Abitbol J, O'Donnell B, Wakefield C, Jewlal E, Kelly J, Barr K, et al. Double deletion of Panx1 and Panx3 affects skin and bone but not hearing. J Mol Med 2019;97(5):723-6.

Binnerts ME, Kim KA, Bright JM, Patel SM, Tran K, Zhou M, et al. R-Spondin1 regulates Wnt signaling by inhibiting internalization of LRP6. Proc Natl Acad Sci U S A 2007;104(37):14700-5.

Churko J, Chan J, Shao Q, Laird D. The G60S Connexin43 Mutant Regulates Hair Growth and Hair Fiber Morphology in a Mouse Model of Human Oculodentodigital Dysplasia. J Invest Dermatol 2011a;131:2197–204.

Churko J, Kelly JJ, MacDonald A, Lee J, Sampson J, Bai D, et al. The G60S Cx43 mutant enhances keratinocyte proliferation and differentiation. Exp Dermatol 2012;21:612-8.

Churko J, Shao Q, Gong X, Swoboda K, Bai D, Sampson J, et al. Human Dermal Fibroblasts Derived from Oculodentodigital Dysplasia Patients Suggest that Patients may have Wound Healing Defects. Hum Mutat 2011b;32(4):456-66.

Fabregat A, Sidiropoulos K, Garapati P, Gillespie M, Hausmann K, Haw R, et al. The Reactome pathway Knowledgebase. Nucleic Acids Res 2016;44(D1):D481-7.

Fabregat A, Sidiropoulos K, Viteri G, Forner O, Marin-Garcia P, Arnau V, et al. Reactome pathway analysis: a high-performance in-memory approach. BMC Bioinformatics 2017;18(1):142.

Freeman TJ, Sayedyahossein S, Johnston D, Sanchez-Pupo RE, O'Donnell B, Huang K, et al. Inhibition of Pannexin 1 Reduces the Tumorigenic Properties of Human Melanoma Cells. Cancers (Basel) 2019;11(102).

Gosset M, Berenbaum F, Thirion S, Jacques C. Primary culture and phenotyping of murine chondrocytes. Nat Protoc 2008;3(8):1253-60.

Hampton AL, Hish GA, Aslam MN, Rothman ED, Bergin IL, Patterson KA, et al. Progression of Ulcerative Dermatitis Lesions in C57BL/6Crl Mice and the Development of a Scoring System for Dermatitis Lesions. J Am Assoc Lab Anim Sci 2012;51(3):586-93.

Holmen SL, Giambernardi TA, Zylstra CR, Buckner-Berghuis BD, Resau JH, Hess JF, et al. Decreased BMD and limb deformities in mice carrying mutations in both Lrp5 and Lrp6. J Bone Miner Res 2004;19(12):2033-40.

Moon PM, Penuela S, Barr K, Khan S, Pin CL, Welch I, et al. Deletion of Panx3 Prevents the Development of Surgically Induced Osteoarthritis. J Mol Med 2015;93(8):845-56.

Mootha VK, Lindgren CM, Eriksson K, Subramanian A, Sihag S, Lehar J, et al. PGC-1α-responsive genes involved in oxidative phosphorylation are coordinately downregulated in human diabetes. Nat Genet 2003;34(3):267-73.

Penuela S, Bhalla R, Gong X-Q, Cowan KN, Celetti SJ, Cowan BJ, et al. Pannexin 1 and pannexin 3 are glycoproteins that exhibit many distinct characteristics from the connexin family of gap junction proteins. J Cell Sci 2007;120(21):3772-83.

Penuela S, Bhalla R, Nag K, Laird D. Glycosylation Regulates Pannexin Intermixing and Cellular Localization. Mol Biol Cell 2009;20:4313–23.

Piprek RP, Kolasa M, Podkowa D, Kloc M, Kubiak JZ. Tissue-specific knockout of E-cadherin (Cdh1) in developing mouse gonads causes germ cells loss. Reproduction 2019;158:147-57.

Press E, Alaga K, Barr K, Shao Q, Bosen F, Willecke K, et al. Disease-linked connexin26 S17F promotes volar skin abnormalities and mild wound healing defects in mice. Cell Death Dis 2017;8:e2845.

Robb KP, Juignet L, Morissette Martin P, Walker JT, Brooks CR, Barreira C, et al. Adipose Stromal Cells Enhance Decellularized Adipose Tissue Remodeling Through Multimodal Mechanisms. Tissue Eng Part A 2021;27(9-10):618-30.

RStudio Team. RStudio: Integrated Development Environment for R. Boston, MA: RStudio, PBC; 2022.

Schindelin J, Arganda-Carreras I, Frise E, Kaynig V, Longair M, Pietzsch T, et al. Fiji: an open-source platform for biological-image analysis. Nat Methods 2012;9:676–82.

Schmidt M, Gutknecht D, Simon J, Schulz J, Eckes B, Anderegg U, et al. Controlling the Balance of Fibroblast Proliferation and Differentiation: Impact of Thy-1. J Invest Dermatol 2015;135:1893–902.

Subramanian A, Tamayo P, Mootha VK, Mukherjee S, Ebert BL, Gillette MA, et al. Gene set enrichment analysis: a knowledge-based approach for interpreting genome-wide expression profiles. Proc Natl Acad Sci U S A 2005;102(43):15545-50.

Szklarczyk D, Gable AL, Lyon D, Junge A, Wyder S, Huerta-Cepas J, et al. STRING v11: protein-protein association networks with increased coverage, supporting functional discovery in genome-wide experimental datasets. Nucleic Acids Res 2019;47(D1):D607-D13.

Szklarczyk D, Gable AL, Nastou KC, Lyon D, Kirsch R, Pyysalo S, et al. The STRING database in 2021: customizable protein-protein networks, and functional characterization of user-uploaded gene/measurement sets. Nucleic Acids Res 2021;49(D1):D605-D12.

Wakefield CB, Lee VR, Johnston D, Boroumand P, Pillon NJ, Sayedyahossein S, et al. Pannexin 3 deletion reduces fat accumulation and inflammation in a sex-specific manner. International Journal of Obesity 2021.

Ye J, Coulouris G, Zaretskaya I, Cutcutache I, Rozen S, Madden TL. Primer-BLAST: A tool to design target-specific primers for polymerase chain reaction. BMC Bioinformatics 2012;13:134.

**Supplementary Table S1. List of genotyping and RT-qPCR primers.**

| Primer | Forward Sequence (5' – 3') | Reverse Sequence (5' – 3') | Reference |
|---|---|---|---|
| *Panx3 R1 (Genotyping)* | CGCAGCATTCCGGAACCTGAG | AACTAGCCGGAGGGGTCG | (Abitbol et al., 2019) |
| *Panx3 RT (Genotyping)* | CGCAGCATTCCGGAACCTGAG | CTTGTCCTGCGTATGGTG | (Abitbol et al., 2019) |
| *Panx3* | TTTCGCCCAGGAGTTCTCATC | CCTGCCTGACACTGAAGTTG | (Abitbol et al., 2019) |
| *Panx1* | ACAGGCTGCCTTTGTGGATTCA | GGGCAGGTACAGGAGTATG | (Abitbol et al., 2019) |
| *Panx2 Full Length* | TGGTACCCATCCTGCTGGT | GGGTGAAGTTGTGCGGAGT | (Abitbol et al., 2019) |
| *Panx2 Isoform 202* | CAGCCCGTGTCTCCTCTCTAC | GTAGCCGCGGGCGTACA | (Abitbol et al., 2019) |
| *Gapdh* | TGCGACTTCAACAGCAACTC | GTAGGCCATGAGGTCCAC | (Abitbol et al., 2019) |
| *Acta2* (α-SMA) | GCTGGACTCTGGAGATGG | GCAGTAGTCACGAAGGAATAG | (Schmidt et al., 2015) |
| *K14* | AGTGAGAAAGTGACCATGCAGA | CTGCCAGGATCTTGCTCTTCAG | (Ye et al., 2012) |
| *K5* | TGAGGAGCTGCAACAGACAG | AGGTTGGCACACTGCTTCTT | (Ye et al., 2012) |
| *Inv* | CTCCTGTGAGTTTGTTTGGTCT | CACACAGTCTTGAGAGGTCCC | (Ye et al., 2012) |
| *Lor* | TCCCTGGTGCTTCAGGGTAAC | TCTTTCCACAACCCACAGGA | (Ye et al., 2012) |
| *K10* | CAGCTGGCCCTGAAACAATC | AGTTGTTGGTACTCGGCGTT | (Ye et al., 2012) |
| *Cdh1* | CCCAAGCACGTATCAGGGTC | GTACACAGCTTTCCACGCC | (Piprek et al., 2019) |
| *Vcl* | TGGTCTAGCAAGGGCAATGA | AAAGCCAGCATCTGTTCGGA | (Ye et al., 2012) |
| *Wnt3* | CAAGCACAACAATGAAGCAGG | TCGGGACTCACGGTGTTTCTC | (Binnerts et al., 2007) |
| *Lrp5* | TGTACTGCAGCTTGGTCCC | CCAGTAAATGTCGGAGTCTACAATG | (Holmen et al., 2004) |
